# Neuronal differentiation and activity drive nucleocytoplasmic shuttling of the intellectual disability kinase TLK2

**DOI:** 10.1101/2025.02.20.639293

**Authors:** Lubna Nuhu-Soso, Heidi Denton, Darren L Goffin, Ines Hahn, Gareth J.O. Evans

## Abstract

Mental retardation autosomal dominant 57 (MRD57) is a rare neurodevelopmental disorder characterised by delayed language and psychomotor development, intellectual disability, hypotonia, gastrointestinal issues and facial dysmorphia. It is linked to genetic mutations in the serine/threonine kinase TLK2, characterised by haploinsufficiency and in some cases, its loss or impaired kinase function. TLK2 is an established cell cycle regulator that has been extensively studied in mitotic cells. It is upregulated in cancers, driving tumour growth, however, the role of TLK2 in postmitotic neurons is not understood. We therefore aimed to gain insight into how TLK2 mutations cause MRD57 by determining where TLK2 is expressed in the brain and its subcellular localisation during neuronal differentiation. Public human and mouse brain transcriptomic data revealed splice variant diversity in the N-terminus of TLK2, which contains its nuclear localisation sequence (NLS). Using splice-specific *in situ* hybridisation probes, we observed expression of TLK2 isoforms that contain and lack the NLS in the mouse hippocampus and cerebellum. We confirmed these findings in human SH-SY5Y neuroblastoma cells, and found that neuronal differentiation of these cells enhances a cytoplasmic pool of TLK2 by two mechanisms: nuclear export of full length TLK2 and increased expression of TLK2 splice variants lacking the NLS. Finally, acute stimuli that mimic synaptic activity were sufficient to elicit nuclear export of TLK2. Our data highlight the need to establish the neuronal cytoplasmic substrates of TLK2 and determine how the loss of TLK2 activity in MRD57 might impact their function in the developing and mature brain.

## Introduction

Tousled-like kinase 1 and 2 are the mammalian homologues of the *Arabidopsis* mutant, *tousled*, which was discovered in a developmental screen (Roe et al., 1993). Indeed, both human TLK1 and TLK2 have been linked with rare developmental disorders (Segura-Bayona and Stracker, 2019; Villamor-Payà et al., 2024). To date, there have been close to 50 documented patients worldwide with TLK2 mutations, classified as autosomal dominant Mental Retardation Disorder 57 (MRD57; (Lelieveld et al., 2016; Reijnders et al., 2018; Töpf et al., 2020; Pavinato et al., 2022; Woods et al., 2022), with a predicted incidence of approximately 3 in 100,000 individuals (López-Rivera et al., 2020). The clinical phenotypes associated with MRD57, like other neurodevelopmental disorders, are heterogeneous, including intellectual disability, delayed psychomotor development in infancy or early childhood, language delay, hypotonia, feeding problems, gastrointestinal issues, dysmorphic facial features, microcephaly, behaviour problems and global developmental delays (Reijnders et al., 2018; Töpf et al., 2020; Pavinato et al., 2022; Woods et al., 2022). The majority of TLK2 mutations in MRD57 are *de novo* mutations, but seven are inherited within four different families (Reijnders et al., 2018; Töpf et al., 2020; Pavinato et al., 2022). Like the *de novo* cases, inherited TLK2 mutations are heterozygous dominant except one report of a homozygous recessive inheritance (Reijnders et al., 2018; Töpf et al., 2020; Pavinato et al., 2022). MRD57 patients have TLK2 mutations throughout the protein and all are predicted to result in a reduction of TLK2 expression or its catalytic activity (Mortuza etc).

Functionally, TLK2 is an established regulator of DNA replication, cell cycle checkpoint recovery and chromatin remodelling (Bruinsma et al., 2016; Kim et al., 2016a; Segura-Bayona et al., 2017; Silljé et al., 1999). TLK2 facilitates cell cycle progression mainly by enhancing histone supply via phosphorylation of its best characterised substrate, ASF1 (Klimovskaia et al., 2014). In-line with other cell cycle kinases, TLK activity and gene copy number are increased in cancer, including breast, cervix, lung, liver, colon and kidney (Bhoir and De Benedetti, 2023; Kim et al., 2016b; Lee et al., 2018). The chromosomal locus of TLK2, 17q23, contains other oncogenes, and is frequently amplified in more than 40% of breast cancer tumours (Kelemen et al., 2009; Kim et al., 2016b). Moreover, single nucleotide polymorphisms (SNPs) in TLK2 have been identified as risk factors for breast cancer (Kelemen et al., 2009). Indeed, inhibition of TLK2 has been identified as a therapeutic strategy and small molecule inhibitors are in development (Kim et al., 2016b; Lee et al., 2018).

In contrast to its role in the cell cycle and cancer, the neuronal expression and cell biology of TLK2 is poorly understood. Here we have characterised its neuronal transcript expression, revealing a conserved splice diversity in rodents and humans. TLK2 alternative splicing occurs in exons encoding the N-terminus, resulting in ‘long’ and ‘short’ isoforms, with the latter lacking the nuclear localisation sequence (NLS). Expression of both long and short TLK2 isoforms is prevalent in neurons of the mouse hippocampus and cerebellum. Using a cell model of neuronal differentiation, we observed the upregulation of short TLK2 expression during neuronal differentiation, with a concomitant shuttling of long TLK from the nucleus to the cytoplasm. Pharmacological treatments mimicking neuronal activity also drive nucleocytoplasmic shuttling of TLK2. These data highlight the likely importance of cytoplasmic TLK2 in terminally differentiated neurons, which have exited the cell cycle. There is hence a need to determine the neuronal substrates of TLK2 to shed light on the neuronal phenotypes of MRD57.

## Materials and Methods

### Materials

SH-SY5Y and Flp-In T-REx SK-N-SH neuroblastoma cell lines were kind gifts from Dr Andrew Stoker, University College London and Dr Han-Jou Chen, University of York respectively. The Flp recombinase expression vector, pOG44 was a kind gift from Dr Paul Pryor, University of York.

### Bioinformatics

TLK2 transcript IDs and exon structures were derived from their Ensembl gene annotations (version 113) using human (GRCh38.p14) or mouse (GRCm39) genome builds (Harrison et al., 2024). The human TLK2 RNAseq data used for the transcriptomic analyses described in this manuscript were obtained from the GTEx Portal, accession number phs000424.v8.p2. Mouse brain TLK2 RNAseq data was obtained from the Human Protein Atlas (proteinatlas.org; (Sjöstedt et al., 2020)). TLK2 transcript expression values in transcripts per million (TPM) were converted to percentages by tissue or brain region.

### RNA extraction, cDNA synthesis and RT PCR

All work involving mice was approved by the University of York Animal Welfare and Ethical Review Body and performed under UK Home Office legislation (project licence P38B2013E). Adult wild-type C57BL/6 mouse forebrain and cerebellum were dissected and homogenised in RIPA buffer (50 mM Tris, pH 8.0, 150 mM NaCl, 1% (v/v) Triton-X-100, 0.5% (w/v) sodium deoxycholate, 0.1% (w/v) SDS, 1 mM EDTA supplemented with 1 mM Na_3_VO_4_, 0.1% (v/v) β-mercaptoethanol, 1 mM PMSF and 1:200 protease inhibitor cocktail (Sigma)) using 5 ml buffer/g of tissue. The homogenate was incubated at 4°C with agitation for 1 h and then centrifuged at 20,000*g* for 30 min at 4°C. RNA was extracted from the supernatant. SH-SY5Y cells undergoing neuronal differentiation were plated at 10^4^ cells/ cm^2^ onto collagen coated 6-well tissue culture dishes. RNA was extracted from the differentiating cells at two day intervals. For both mouse brain and SH-SY5Ys, RNA was extracted using a NucleoSpin RNA isolation kit (Macherey-Nagel) according to the manufacturer’s instructions. Complementary DNA was synthesised from 1 µg of RNA using a SuperScript IV First-Strand cDNA Synthesis Kit (Life Technologies) according to the manufacturer’s instructions in the presence of both oligo d(T)_18_ and random hexamer primers. A no RT cDNA sample was performed for each time point and cDNA was stored at -20°C. PCR reactions were performed in 25 µl reactions containing 1 µl of cDNA, 1X GoTaq flexi buffer, 0.2 mM each of dNTP mix, 0.4 µM of each primer pair (Table 1) and 1U Taq polymerase. Cycling conditions consisted of an initial denaturing step at 98°C for 30s followed by 25-30 cycles of 95°C for 10s, 55°C for 30s, 72°C for 1 min/kb and a final extension at 68°C for 10 min. 5 - 10 µl of the PCR products were separated on 1.5-2% (w/v) agarose gels to confirm amplification.

**Table 1.**
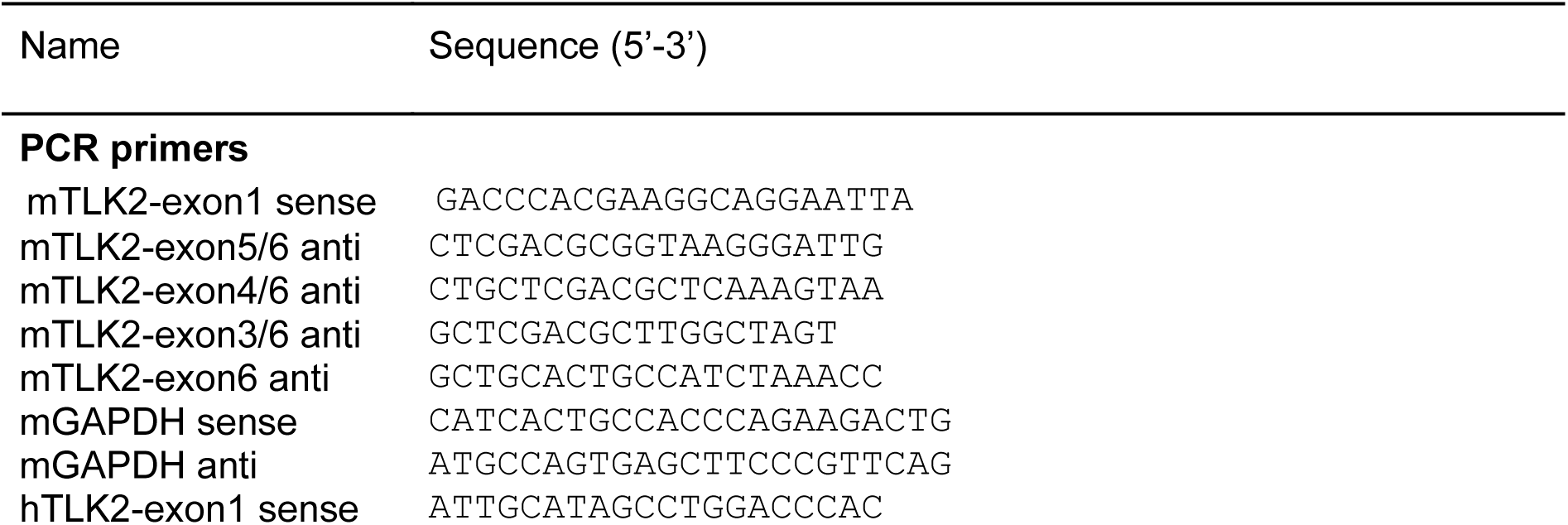
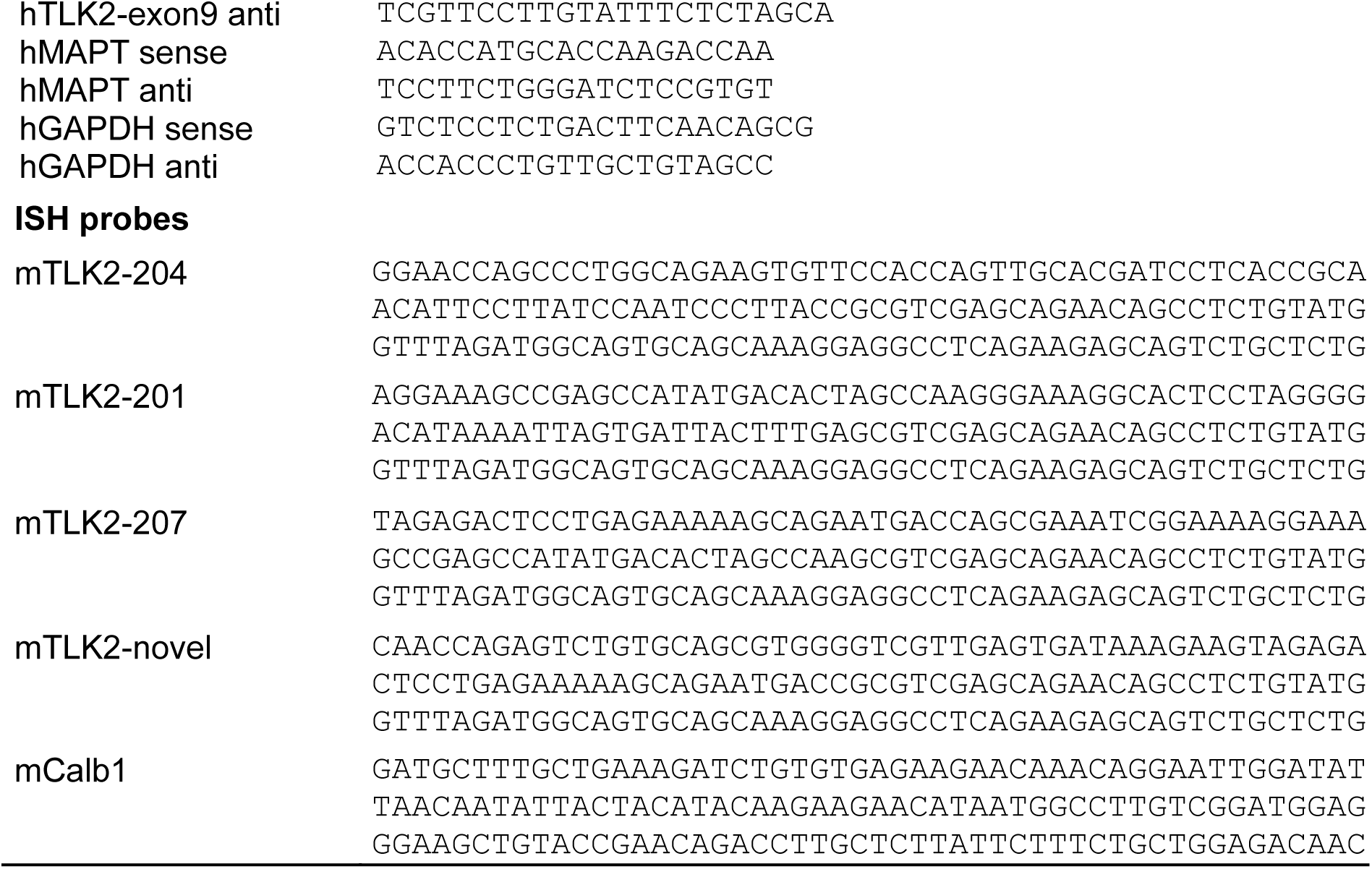
Primers and probes used in this study.

### *In situ* hybridisation

Mice were anaesthetised with isoflurane inhalation followed by an intraperitoneal injection of 100 mg/kg ketamine and 12.5 mg/kg xylazine and then transcardially perfused with 4% paraformaldehyde (PFA) in 0.1 M sodium-potassium PBS. The brain dissected and post-fixed in 4% PFA overnight at 4°C. Following a further 24 h incubation in 30% sucrose, brains were embedded in OCT, frozen at -80°C, and 25 μm coronal or sagittal sections were cut using a Leica CM1950 cryostat. Probes for *in situ* hybridisation were transcribed using 10X digoxigenin (DIG) RNA labelling mix (Roche) and T7 polymerase. For *in vitro* transcription, template DNA was synthesised as gBlocks™ (IDT) and PCR amplified to introduce the T7 promoter sequence (Table 1). *In situ* hybridisation was performed according to (Fisher et al., 2002) with adaptations for sections mounted on slides. Briefly, the sections were first incubated in 4% PFA for 10 min at room temperature (RT) followed by two 5 min PBST washes. Sections were then treated with 4 µg/ml proteinase K (Roche) in PBST for 10 min and then washed twice in PBST for 5 min. A second fix in 4% (w/v) PFA was performed for 10 min, followed by two 5 min PBST washes. Next, the sections were acetylated using 0.5% (v/v) acetic anhydride in 0.1 M triethanolamine, pH 7.8 for 10 min followed by two 5 min PBST washes and finally a 15 min wash in 5X saline-sodium citrate buffer (SSC; Sigma). Pre-hybridisation was carried out by incubating the sections in hybridisation buffer (50% (v/v) formamide (Ambion, AM9342), 5X SSC, 100 µg/ml heparin, 1X Denhardt’s, 0.1% (v/v) Tween-20, 0.1% (w/v) CHAPS, 10 mM EDTA, 1 mg/ml total yeast RNA) containing no probe at 65°C for 2-5 h in a humidified box. Hybridisation followed by incubating the sections with 300 ng/ml of probe solution (DIG prep in hybridisation buffer) overnight at 65°C in a humidified box.

Post-hybridisation, the sections were subjected to several hot wash steps. First a 10 min wash in hybridisation buffer containing no probe at 65°C, then a 1 h wash in 2X SSC followed by a 1 h wash in 0.2X SSC, both at 65°C. The sections were then washed twice for 5 min in MABT (100 mM maleic acid, 150 mM NaCl, 0.1% (v/v) Tween-20, pH 7.8) at RT and then blocked in blocking solution (2% (w/v) Boehringer Mannheim Blocking Reagent (Roche, 11096176001), 20% (v/v) heat-treated lamb serum (Sigma), 1X MAB) for 2 h at RT in a humidified box. Next, the sections were incubated with anti-DIG-AP (Table 2) in blocking solution overnight at 4°C in a humidified box. Post-antibody washes were carried out at RT by washing the sections three times for 1 h in MABT and once in alkaline phosphatase (AP) buffer (100 mM Tris, pH 9.5, 50 mM MgCl_2_, 100 mM NaCl) for 10 min. The colour reaction was initiated by adding a 1:3 dilution of BM purple (Roche, 11442074001) in AP buffer to the sections and incubation at RT in a humidified box for 1-5 days. The reaction was stopped by washing the sections twice for 10 min in PBST, fixing in 4% (w/v) PFA for 1 h or overnight at 4°C and a 15 min wash in PBST. The sections were air dried, mounted in aqueous mounting media (Sigma, 324590) and stored at 4°C. Images were acquired on a Leica DM2500 microscope with a 10X objective (sagittal) and a Zeiss Stemi 508 stereo microscope (coronal).

**Table 2.**
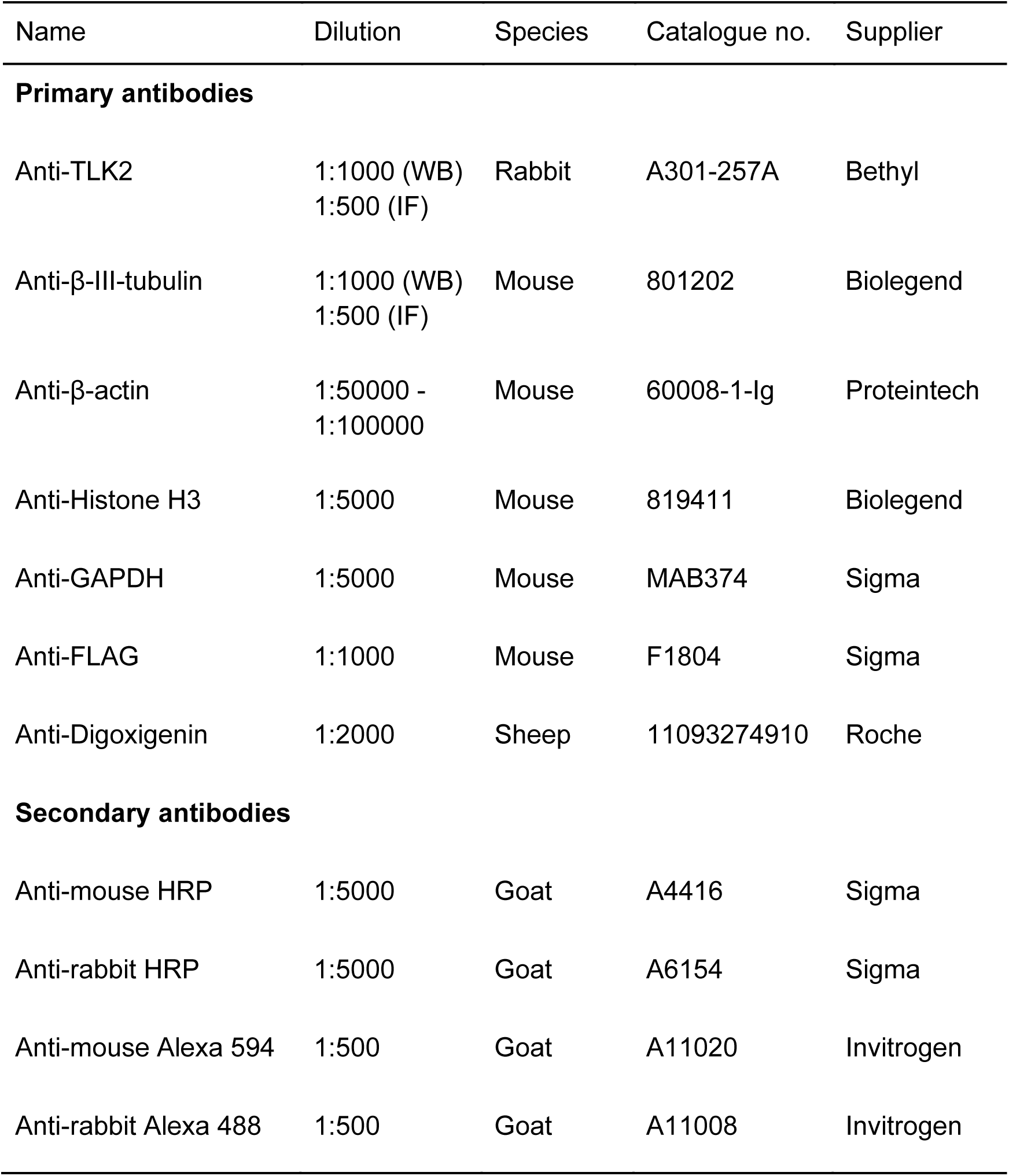
Antibodies used in this study.

### Culture, neuronal differentiation and pharmacological treatment of SH-SY5Y cells

SH-SY5Y cells were maintained at 37°C, 5% CO_2_ in a humidified atmosphere in Dulbecco’s Modified Eagle Medium (DMEM) with high glucose, pyruvate and L-glutamine supplemented with 10% foetal bovine serum (FBS) and 1% penicillin/streptomycin.

Neuronal differentiation of SH-SY5Y cells was performed according to a previously described method (Encinas et al., 2000). Briefly, tissue culture dishes or coverslips were coated in 0.05 mg/ml collagen in 60% (v/v) ethanol. Cells were plated at 10^4^ cells/cm^2^ (for protein analysis) or 5 x 10^3^ cells/well (for microscopy) in DMEM supplemented with 15% (v/v) heat inactivated FBS and 1% (v/v) penicillin/streptomycin. Twenty four hours after plating, the media was changed and supplemented with 10 µM retinoic acid (RA; Sigma, R2625). Six days after plating, the cells were washed three times in serum-free DMEM and then incubated with serum-free DMEM containing 50 ng/ml brain-derived neurotrophic factor (BDNF; Qkine) and 1% (v/v) penicillin/streptomycin and cultured for a further seven days. Throughout the differentiation process, the media was changed every two days and supplemented with the appropriate differentiation agent.

For analysing protein expression in differentiating cells, lysates were collected at two day intervals. Cells were washed twice in sterile ice-cold PBS and scraped in 100 µl of lysis buffer (20 mM HEPES, pH 7.4, 10 mM KCl, 2 mM MgCl_2_, 1 mM EDTA and 1 mM EGTA, 1 mM DTT and protease inhibitor cocktail). Lysates were incubated on ice for 15 min, passed through a 25G needle ten times and protein concentration was determined by Bradford assay. The lysates were diluted in Laemmli sample buffer, boiled for 10 min at 90°C and 10 µg of protein was separated by SDS-PAGE and analysed by Western blotting.

Pharmacological stimulation of undifferentiated SH-SY5Y cells was performed on cells plated 24 h prior to treatment on 13 mm coverslips in a 24-well plate at 5 x 10^4^ cells/well. Cells were washed twice in PBS and then incubated for 1 h at 37°C, 5% CO_2_ in either control buffer (170 mM NaCl, 3.5 mM KCl, 0.4 mM KH_2_PO4, 10 mM HEPES, 5 mM NaHCO_3_, 5 mM glucose, 1.2 mM Na_2_SO_4_, 1.2 mM MgCl_2_, 1.3 mM CaCl_2_, pH 7.4, 0.2% (v/v) DMSO) or KCl/FSK buffer (control buffer with 73.5 mM NaCl, 100 mM KCl, 10 µM forskolin, 100 µM IBMX, 0.2% (v/v) DMSO). Cells were then fixed and immunocytochemistry was performed as described below using anti-TLK2.

### Inducible stable TLK2 cell lines

Tetracycline-inducible SK-N-SH cell lines expressing either full length TLK2-203 or short N-terminally truncated TLK2-213 were generated using the Flp-In™ T-REx™ System (Invitrogen). pcDNA5/FRT/TO-FLAG-TLK2 was purchased from the MRC PPU Reagents and Services facility (College of Life Sciences, University of Dundee). TLK2-213 was made by subcloning its ORF from full length TLK2 and ligating it back into the same backbone via 5’ and 3’ NotI restriction sites with the N-terminal FLAG tag intact. Cells were co-transfected with the FRT backbone containing the gene of interest (TLK2-203 or TLK2-213) and a Flp-recombinase expression vector (pOG44). Positive clones were selected by treating with 0.5 mg/ml hygromycin B and assaying for protein expression after the addition of 1 µg/ml doxycycline.

### Subcellular fractionation

Flp-In SK-N-SH cells were fractionated according to the Abcam subcellular fractionation protocol. TLK2 expression was induced using 1 µg/ml doxycycline and left in culture for 48 h prior to fractionation. Briefly, cells were scraped in 500 µl of lysis buffer (described above) and incubated on ice for 15 min. The scraped cells were passed through a 25G needle ten times, incubated on ice for 20 min and then centrifuged at 720*g* for 5 min at 4°C. The post-nuclear supernatant was transferred to a fresh tube. The nuclear pellet was washed by first resuspending in lysis buffer, passing the sample through a 25G needle ten times and centrifuging at 720*g* for 10 min at 4°C. The washed pellet was resuspended in TBS containing 0.1% (w/v) SDS, and sonicated on ice for 3 s. The protein concentration of each fraction was determined by Bradford assay and 20 µg of each fraction was separated by SDS-PAGE and analysed by Western blotting.

### Western blotting

Proteins separated by SDS-PAGE (7.5-12%) gels were transferred onto PVDF membranes (Millipore). Membranes were incubated with primary antibodies (Table 2) overnight at 4°C with agitation. Membranes were washed and incubated with appropriate HRP-conjugated secondary antibodies and immunoreactivity was visualised by incubation with enhanced chemiluminescence reagent and imaged on an iBright FL1000 scanner (Invitrogen).

### Immunocytochemistry and fluorescence microscopy

Cells were washed 3x in PBS and then fixed in 4% (w/v) paraformaldehyde (PFA) in PBS for 20 min at RT. Next cells were washed 3x in PBS then permeabilised and blocked in PBS containing 0.1% (v/v) TritonX-100 and 1% (w/v) bovine serum albumin (BSA). Primary antibodies were diluted in PBS containing 1% (w/v) BSA at appropriate dilutions (Table 2) and incubated on cells overnight at 4°C. Cells were washed 3x in PBS then incubated with secondary antibodies in PBS containing 1% (w/v) BSA for 1 h in the dark at RT. Finally, cells were washed 3x in PBS, 1x in dH_2_O, air dried and mounted in Mowial containing 1 µg/ml 4′,6-diamidino-2-phenylindole (DAPI). Images were acquired in the Bioscience Technology Facility (University of York) on a Zeiss LSM 880 confocal microscope on a 40x/1.4 Oil DIC III objective.

### Data analysis

Quantification of hippocampal *in situ* hybridization images was performed on 16 bit grayscale images in FIJI (Schindelin et al., 2012). The oval selection tool was used to draw regions of interest (ROI) in different locations within the hippocampal formation (CA1, CA3, DGs and DGi). The intensity was then measured and normalised to DGs for each section and probe. Three to four hippocampi were quantified for each probe and the values were plotted as a heatmap in Morpheus (https://software.broadinstitute.org/morpheus).

The quantification of cytoplasmic/nuclear TLK2 ratios of immunofluorescent images was conducted by measuring the TLK2 intensity in the nucleus and cytoplasm of the cells in FIJI using the ROI 1-click plug-in (Thomas and Gehrig 2020). Three hundred cells for each condition were analysed from three technical replicates (100 cells/replicate). Western blots were quantified by densitometry analysis in FIJI. Statistical significance was assessed by t-test or non-parametric ANOVA with Dunn’s post-hoc test. Statistical analyses and graph plotting was performed in RStudio (RStudio Team, 2020).

## Results

### TLK2 is alternatively spliced in human and mouse brain

Since the neuronal distribution and function of TLK2 has not been investigated, we first sought to characterise TLK2 transcript expression in the brain. Previous studies have observed tissue-specific transcripts of TLK2, in the mouse testis for example (Shalom and Don, 1999), and TLK2 immunoblots often have multiple bands that vary between tissues and cell lines (Kim et al., 2016b; Segura-Bayona et al., 2017). Such bands could result from post-translational modifications or alternative splicing of TLK2 or a combination. We began by interrogating publicly available human (GTEx) and mouse brain (Human Protein Atlas) transcriptomic data to characterise neuronal TLK2 transcript expression (Figure 1A,B), focussing on the hippocampus and cerebellum. In the mouse brain, a full length transcript corresponding to the canonical Ensembl TLK2 transcript was the most abundant transcript (mTLK2-204; mouse TLK2 transcripts will be referred to here as ‘mTLK2’). In the human brain, surprisingly the most abundant transcript was not canonical TLK2 (TLK2-203), but a transcript lacking exon 5 (TLK2-202; Figs. 1B, S1 & S2). We also identified further evolutionarily conserved splice variant diversity at the N-terminus of the protein, encoded by exons 1-9. Not surprisingly, there were fewer alternative splicing events in exons encoding the middle coiled-coil domain and C-terminal kinase domain of TLK2, which are essential for dimerisation and catalytic activity respectively (Mortuza et al., 2018); Figure 1B). Interestingly, splice variants lacking exon 3 in which the nuclear localisation sequence (NLS, Figure 1A) is located (Yamakawa et al., 1997) are also abundant (Figure 1B,S1). This suggests that a pool of TLK2 is constitutively resident outside the nucleus. Henceforth in this study we shall refer to isoforms that include the NLS as ‘long TLK2’, and those lacking the NLS as ‘short TLK2’. In the human GTEx dataset it was observed that transcript TLK2-207 represents nearly a third of TLK2 mRNA expression, but is predicted to be subject to nonsense mediated decay due to an intron retention at exon 21 (Figure 1B).

**Figure 1.**
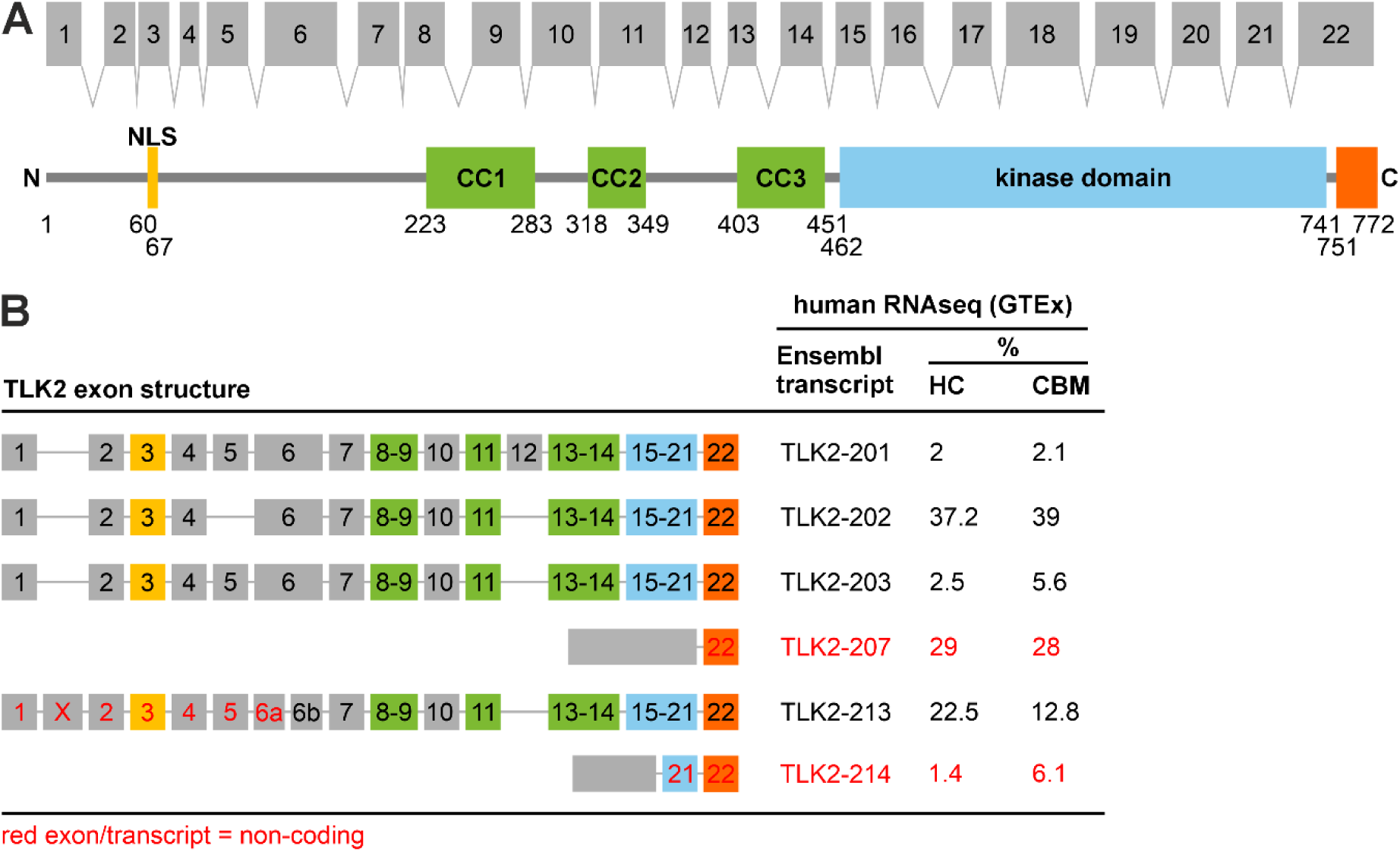
TLK2 has splice variant diversity in the human brain. (A) Exonic structure of human TLK2, mapped onto a schematic representation of its protein domains. NLS, nuclear localisation sequence; CC, coiled-coil domain. (B) Table of % TLK2 transcript abundance in the human hippocampus and cerebellum derived from GTEx RNAseq data. Exons and transcripts labelled in red are predicted to be non-protein coding. Only transcripts present >2% in either brain region are included.

To confirm the existence of alternative N-terminal TLK2 splice variants in the brain, we performed RT-PCR of mouse brain TLK2 mRNA. The annotated transcripts of mouse TLK2 have splice diversity within exons 1-6 (Figure 2A and Figure S2). We first amplified and sequenced all N-terminal splice variants in mouse forebrain or cerebellar cDNA using transcript-specific primers and also a pair of ‘pan isoform’ primers located in exon 1 (forward) and exon 6 (reverse), which yielded four bands (Figure 2A). Three of the bands corresponded to the 5’ of annotated isoforms, two of which are predicted to contain the NLS (mTLK2-204 and -201) and one that lacks the NLS due to an alternative start site in exon 6 (mTLK2-207). We also sequenced a novel isoform that has a truncated exon 3, lacks exon 4 and 5 and would also not encode the NLS (mTLK2-novel; Figure 2B and Figure S2). We next used *in situ* hybridisation to observe whether there is any cell type specificity in the expression of these TLK2 isoforms in the brain. All mouse TLK2 transcripts express exon 6 and combinations of exons 1-5, therefore we designed a suite of short 150 bp splice junction-spanning *in situ* probes anchored at the 3’ end by the first 75 bp complementary to exon 6 and containing a variable 5’ 75 bp complementary to exons 5, 4 or 3 (Figure 2B).

**Figure 2.**
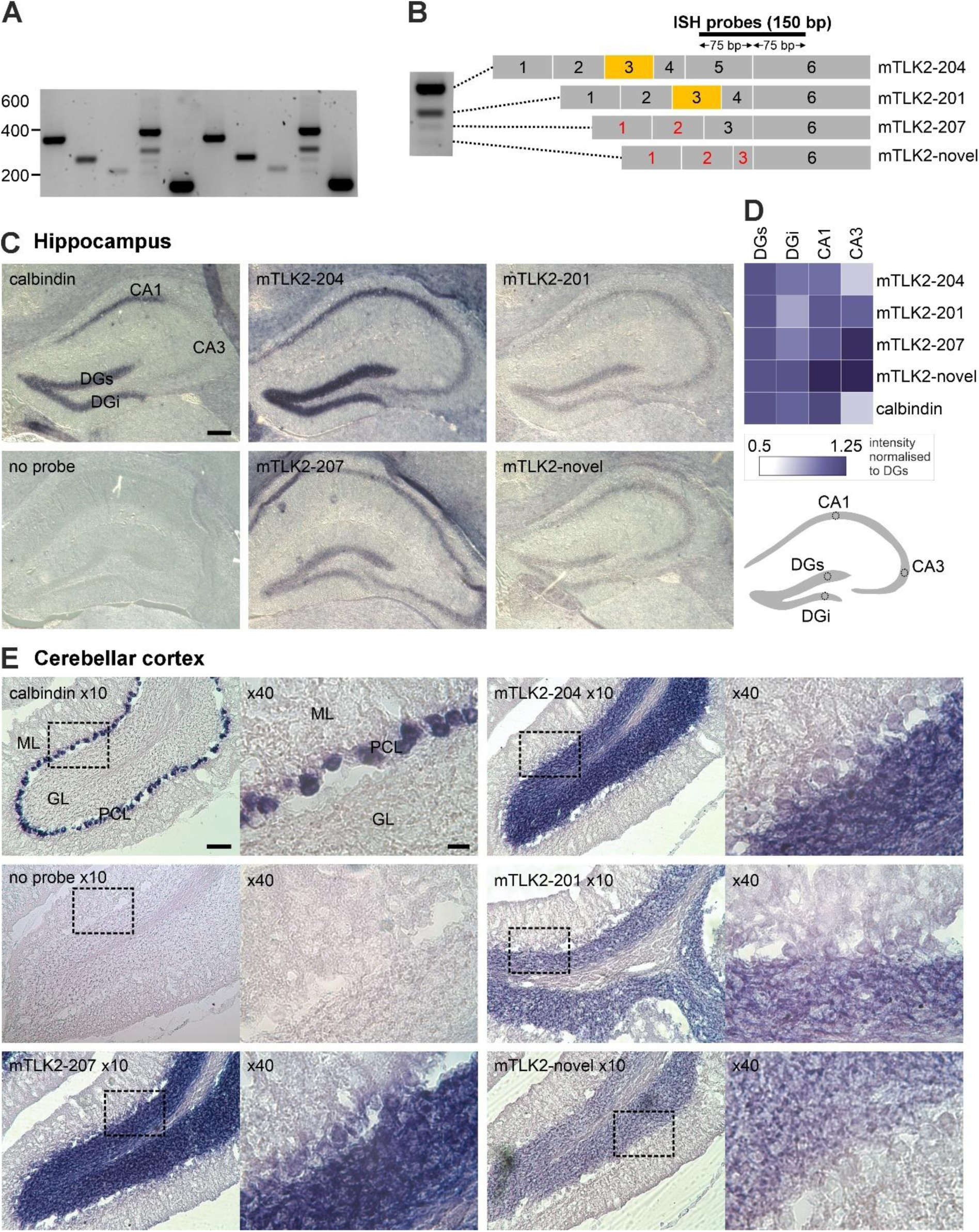
Multiple isoforms of TLK2 are expressed in mouse forebrain and cerebellum. (A) RT-PCR of TLK2 splice variants from mouse forebrain and cerebellum cDNA. Transcript specific primer pairs or pan exon 1-6 primers. (B) Left, lane from PCR of exon 1-6 primer pair. The four bands were excised and sequenced. Right, exon structure of sequenced mouse brain splice variants, and location of *in situ* hybridisation probes. *In situ* hybridisation was performed on 25 µm frozen sections from coronal (C) or sagittal (E) P56 mouse brain sections. The indicated 150 bp exon junction spanning probes were used to examine calbindin or TLK2 transcript expression in the hippocampus (C) or cerebellar cortex (E). Representative images are shown from n=3 sections from two mouse brains. Cerebellar cortex images are shown at x10 and x40 magnification, indicated by the insert box. DG, dentate gyrus; ML, molecular layer; GL, granule layer; PCL, Purkinje cell layer. Scale bars: hippocampus 100 µm, cerebellar cortex x10, 200 µm; x40, 20 µm. (D) Heatmap depicting quantification of TLK2 splice variant expression within the dentate gyrus, CA1 and CA3 sub-regions of the hippocampal formation. For each section, pixel intensity was measured in regions of interest (ROI), as indicated in the cartoon and normalised to the ROI located in the suprapyramidal blade of the dentate gyrus. Three sections were quantified for each probe.

In coronal and sagittal mouse brain sections we examined the expression of TLK2 isoforms in the hippocampal formation and cerebellar cortex respectively. A 150 bp exon junction spanning calbindin probe was used as a positive control, confirming calbindin expression in the Purkinje neurons of the cerebellar cortex and dentate gyrus and CA1 neurons of the hippocampus (Figure 2C,E). Low background staining was observed in the absence of an *in situ* probe (Figure 2C,E). In the coronal sections, hippocampal staining of all TLK2 isoform probes was detected in excitatory neurons of the dentate gyrus, CA1 and CA3 (Figure 2C). Although quantifying the intensity of staining between probes was not appropriate, using regions of interest placed in the DG, CA1 and CA3 fields, we quantified the relative intensity of staining of each TLK2 isoform in these hippocampal cell types and normalised the density to the signal in the suprapyramidal blade of the dentate gyrus (DGs; Figure 2C). The full length TLK2 (exons 1-6) probe, corresponding to the most abundant splice variant mTLK2-204, gave strong staining in the cell soma of the granule neurons in the DG and pyramidal neurons in CA1 and to a lesser extent in CA3 (Fig 2B, C). The other long TLK2 isoform (mTLK2-201) was less readily detected in the hippocampus, but quantification revealed differential expression in the two pyramidal blades of the DG, with the suprapyramidal blade having the highest intensity (Figure 2D). The relative staining of the shorter variant, mTLK2-207, was highest in CA1 neurons. In-line with the PCR (Figure 2A), staining of the novel short TLK2 isoform appeared weakest, with the highest intensity in CA1 and CA3 compared to the DG (Figure 2C, D).

In sagittal sections of the cerebellar cortex, staining of the TLK2 probes was readily detectable in the abundant granule neurons and to a lesser extent in Purkinje neurons (Figure 2E). The intensity of staining for the full length mTLK2-204 probe was similar to that of the shorter mTLK2-207 in the granule cell and Purkinje layers, whereas the mTLK2-201 and mTLK2-novel probes had weak granule layer staining and barely detectable staining of Purkinje neurons (Figure 2E). For all TLK2 probes, we failed to observe staining of inhibitory interneurons in the hippocampus or cerebellar cortex. Taken together, these data confirm the RNAseq data that alternative splice variants of TLK2 are expressed in neurons of the mammalian brain. It is currently unclear as to the physiological relevance of the neuronal cell-type specific expression of these TLK2 variants we have observed.

### Alternative splicing of TLK2 is regulated during neuronal differentiation

To date, our knowledge of TLK2 is mainly limited to its nuclear functions (Segura-Bayona and Stracker, 2019). The possibility of a non-nuclear pool of TLK2 in post-mitotic neurons has interesting implications for the role of TLK2 in the developing and mature brain. To study TLK2 expression in neuronal differentiation we adopted a protocol for differentiating the human neuroblastoma cell line SH-SY5Y. Based on a previously published method we used the sequential application of retinoic acid (RA) and then BDNF with serum withdrawal over several days (Encinas et al., 2000), which produced robust neuronal morphology (Figure 3A) and enhanced transcript expression of the neuronal marker tau (Figure 3B). To capture the diversity of 5’ TLK2 splice variants we performed RT-PCR on SH-SY5Y cDNA samples from day 1-13 with primers located in exons 1 (forward) and 9 (reverse). This yielded six bands (Figure 3B), which were sequenced (Figure S3). Figure 3B summarises the exon structure of each TLK2 PCR product, any corresponding annotated Ensembl transcripts and the predicted mass of the protein they encode. The three longest PCR products mapped onto several Ensembl transcripts with alternative splicing of combinations of exons 5, 7 and 8. The top two bands each potentially correspond to two Ensmbl isoforms due to alternative splicing at exon 12, which wasn’t amplified in the PCR reaction. These bands were also the most prominent PCR products in HEK293 cells. Sequencing the third band yielded a mix of sequences that we were unable to deconvolve, but likely comprises up to three isoforms of the same mass (Figure 3B, S3). We were not able to detect expression of a transcript corresponding to TLK2-213, the annotated TLK2 transcript that lacks the NLS (Figure 1B). Instead, the three smallest PCR products represented novel isoforms lacking exons 3, 4 and 5, with alternative splicing of exons 2, 7 and 8 (Figure 3B). All three of these transcripts are predicted to be protein coding and lack the NLS. HEK293 cells possessed very low levels of these PCR products.

**Figure 3.**
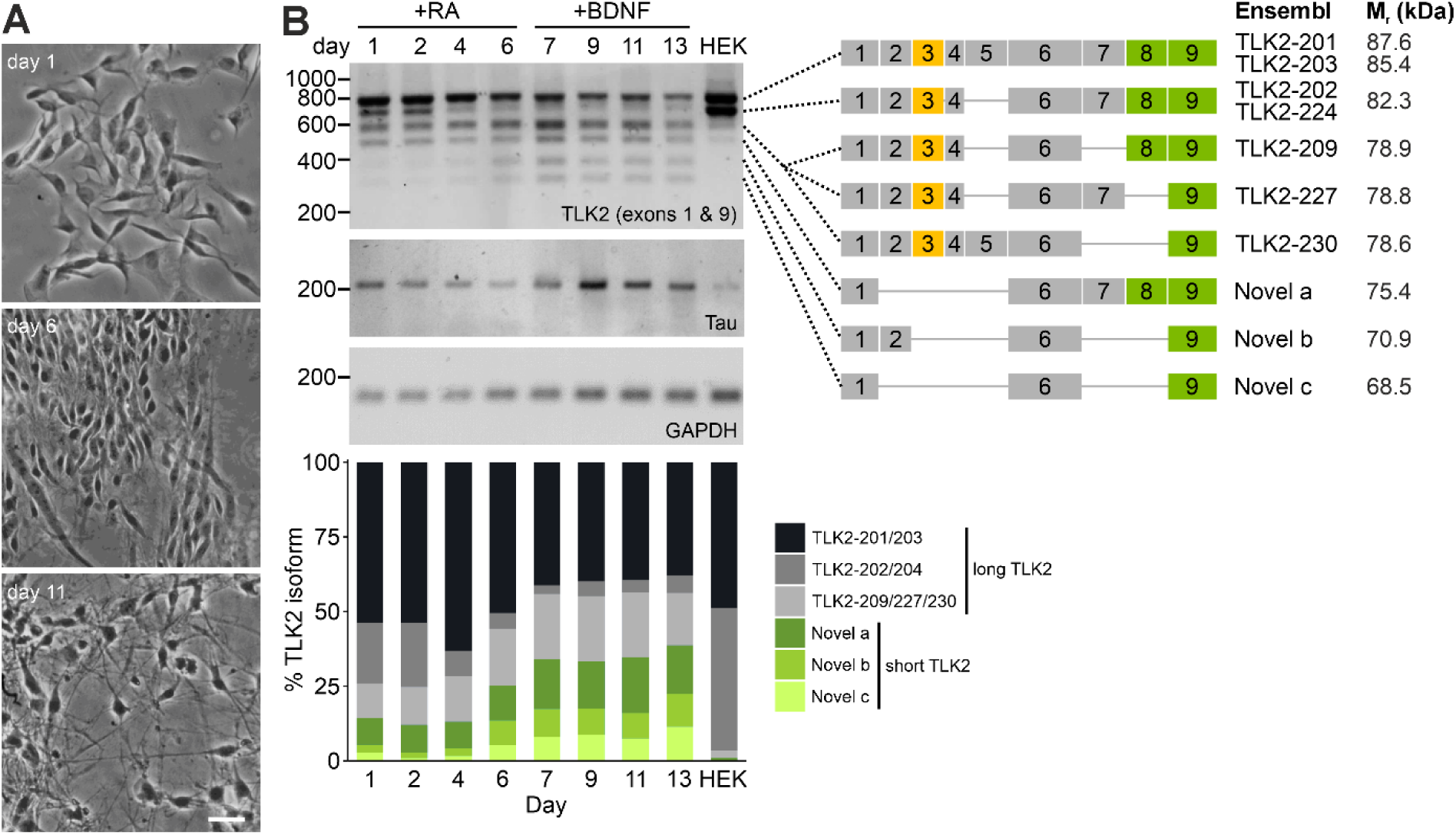
Alternative splicing of TLK2 is regulated during neuronal differentiation. SH-SY5Y cultures were differentiated by sequential treatment with retinoic acid (RA) and brain derived neurotrophic factor (BDNF) as described in Materials and Methods. (A) Brightfield images of SH-SY5Y cultures at the indicated days of differentiation. Scale bar = 40 μm. (B) cDNA was prepared from SH-SY5Y cells at the indicated time points, and HEK293 cells (control) and subjected to PCR with primers for exons 1 and 9 of human TLK2, Tau, and GAPDH. The six TLK2 bands obtained from the cells were excised and subjected to Sanger sequencing. The right hand schematic summarises the exon structure of each sequence and any corresponding Ensembl transcripts that these align to and the predicted molecular mass of the encoded protein. The bottom plot presents quantitative densitometry of each TLK2 RT-PCR product, expressed as a percentage of the total intensity at each timepoint.

We observed a striking regulation of TLK2 isoform expression during neuronal differentiation of the SH-SY5Y cells (Figure 3B). In undifferentiated cells the TLK2 isoforms are dominated by the long transcripts (TLK-201/203; TLK-202/224; TLK2-209/227/230) and the longest of the short isoforms that lacks the NLS (TLK2-novel-a). After the addition of RA, the expression of the long TLK-202/224 isoform is rapidly downregulated (Figure 3B,C). The subsequent addition of BDNF in serum-free media induced an upregulation of the shortest variants (TLK2-novel-b and -c) and a concomitant downregulation of the longer isoforms. At the end of the differentiation protocol, the TLK2 isoform profile was a broader mix of long and short variants with similar levels of expression (Figure 3B).

To establish the corresponding pattern of TLK2 isoform protein expression during differentiation, SH-SY5Y whole cell lysates from the timecourse experiment were subjected to immunoblotting with a C-terminal reactive antibody, for which the epitope is predicted to be present in all major splice variants of TLK2 (Figure 4A). Based on the predicted molecular weight of the proteins encoded by the TLK2 transcripts identified in SH-SY5Y cells (see Figure 3B), we expected that the long isoforms would migrate in the range 78-87 kDa, while the shorter isoforms lacking the NLS would migrate at 68-75 kDa. Indeed, two groups of bands were observed at approximately 85 kDa and 70 kDa, and their expression was regulated during SH-SY5Y differentiation, broadly correlating with the expression of the long and short TLK2-transcripts observed in Figure 3. Quantification of replicate immunoblots by densitometry revealed that the 85 kDa band had a biphasic expression during differentiation, which reduced during RA treatment and then stabilised during BDNF treatment (Figure 4A,B). The ∼70 kDa band increased in expression during differentiation, with this band becoming the dominant isoform in the fully differentiated cells (Figure 4A,B). Taken together these data suggest that neuronal differentiation enhances expression of a TLK2 isoform that lacks the NLS and is predicted to be cytoplasmic.

**Figure 4.**
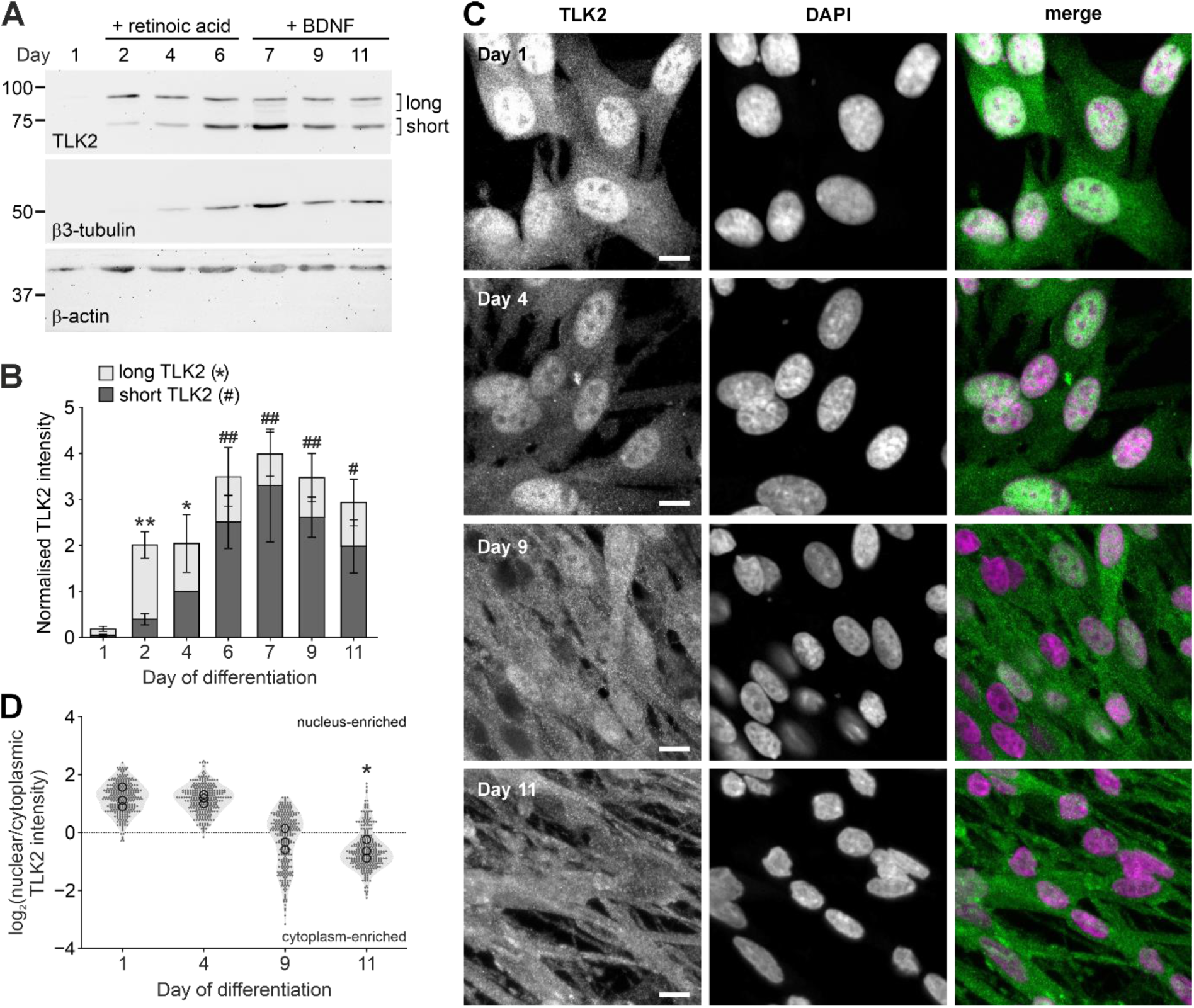
Neuronal differentiation alters the nuclear-cytoplasmic distribution of TLK2. SH-SY5Y cells were lysed or fixed at the indicated timepoints during neuronal differentiation and subjected to immunoblotting with anti-TLK2, anti-β3-tubulin or anti-actin (A) or immunocytochemistry with anti-TLK2 (1:1000; green) and DAPI staining (purple) Scale bar = 10 μm. (C). (B) Quantification of TLK2 immunoblot band intensity from (A), data plotted as stacked bars and raw data points for long TLK2 (yellow circles) and short TLK2 (green circles) isoforms. Data were normalised to the intensity of actin and then day 4 short TLK2 (n=3 independent experiments). (D) TLK2 intensity was measured in regions of interest in the nucleus and cytoplasm of 100 cells from each timepoint (n=3 independent experiments). Data are plotted as log_2_(nuclear/cytoplasmic) TLK2 intensity at each timepoint (grey filled circles). Hollow black circles represent the mean of each replicate. For (B) and (D), data were analysed by Kruskal-Wallis two-tailed ANOVA and Dunn’s post-hoc test; * p < 0.05; ** p < 0.01 compared to day 1. For significance in (B), * = long TLK2 and # = short TLK2.

### Cytoplasmic localisation of TLK2 in differentiated neurons

Immunofluorescent staining of SH-SY5Y cells at 1, 4, 9 and 11 days of differentiation with the TLK2 C-terminal antiserum, revealed a time-dependent relocalisation of TLK2 from the nucleus to the cytoplasm (Figure 4C). We quantified its subcellular localisation by measuring the intensity of TLK2 staining in regions of interest within the cytoplasm and nucleus of the cells. The plot of log2(cytoplasm/nucleus) in Figure 4D revealed that TLK2 is significantly enriched in the cytoplasm at day 11 compared to day 1.

To confirm that TLK2 proteins encoded by splice variants lacking the NLS-containing exon 3 localise to the cytoplasm, we generated stable tet-inducible SK-N-SH T-REx-Flp-in cell lines that express either FLAG-TLK2(180-750), corresponding to the predicted protein encoded by Ensembl isoform TLK2-213 or full length FLAG-TLK2 corresponding to the protein product of the canonical Ensembl TLK2 full length isoform (TLK2-203). Following a 48 h induction of TLK2 expression with doxycycline in these cell lines, we performed a crude subcellular fractionation (Figure 5B). The cells were subjected to hypotonic lysis and we isolated the pellet (enriched in nuclei) and post-nuclear supernatant. Immunoblotting with antibodies raised to compartment markers, histone H3 and GAPDH, confirmed enrichment of nuclear and cytoplasmic proteins respectively (Figure 5B). The immunostaining of anti-FLAG of the subcellular fractions and in fixed cells demonstrated that the N-terminal truncated kinase was exclusively localised to the cytoplasm, while the full length kinase is predominantly nuclear (Figure 5A,B). Taken together we propose that a cytoplasmic pool of TLK2 increases during neuronal differentiation due to two phenomena: an increase in the expression of a short TLK2 splice variant lacking the NLS and the export of full length TLK2 from the nucleus.

**Figure 5.**
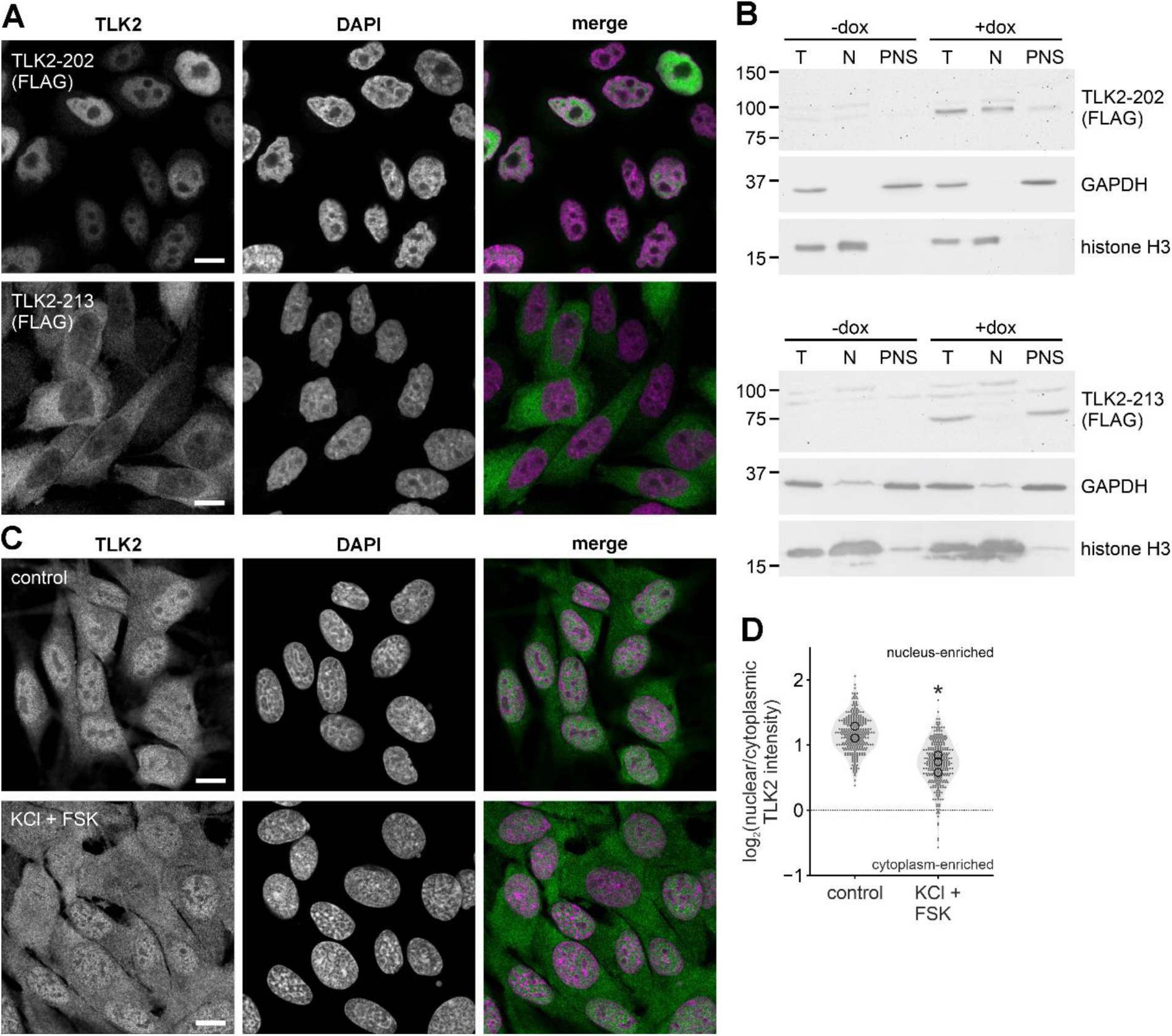
Activity-dependent redistribution of TLK2. Inducible Flp-in SK-N-SH cells expressing FLAG-tagged long TLK2-203 or short TLK2-213 were treated with or without 1 μg/ml doxycycline (dox) for 48 h. (A) Representative immunofluorescence images of dox-treated cells stained with anti-FLAG (1:1000) and DAPI. Scale bar = 10 μm. (B) Cells plated in dishes were lysed and subject to subcellular fractionation to yield a nuclear pellet and postnuclear supernatant. The samples were subjected to immunoblotting with anti-FLAG, anti-GAPDH and anti-histone H3. (C) SH-SY5Y cells equilibrated in KREBS buffer were incubated for 1 h in KREBS buffer with or without (control) 100 mM KCl and 10 μM forskolin (FSK). Fixed cells were stained with anti-TLK2 and DAPI. Scale bar = 10 μm. (D) TLK2 intensity was measured in regions of interest in the nucleus and cytoplasm of 100 cells from each treatment (n=3 experiments). Data are plotted as log_2_(nuclear/cytoplasmic) TLK2 intensity at each timepoint (grey filled circles). Hollow black circles represent the mean of each replicate and were analysed by t-test; * p < 0.05 compared to control.

### Nuclear-cytoplasmic shuttling of TLK2

The nuclear-cytoplasmic transport of TLK2 has been previously reported by Zhang et al (1999) who discovered a yeast 2-hybrid interaction between a 14-3-3**ζ** bait and a C-terminal clone of TLK2 (amino acids 657-750). They found that 14-3-3**ζ** immunoprecipitated from cell lysates with TLK2 and a dominant negative 14-3-3 promoted the nuclear localisation of TLK2 (Zhang et al., 1999). Since 14-3-3 proteins act by binding a phosphorylated motif (Yaffe et al., 1997), we hypothesised that during neuronal differentiation, phosphorylation of TLK2 recruits 14-3-3, thus preventing nuclear import of TLK2 and restricting it to the cytoplasm. Using undifferentiated SH-SY5Y cells, we attempted to mimic these signals to drive cytoplasmic localisation of endogenous TLK2 by stimulating kinase signalling with 100 mM KCl and 10 uM forskolin for 1 h. 100 mM KCl is known to depolarise SH-SY5Y cells, mimicking synaptic activity, and should activate Ca^2+^-dependent intracellular signalling in addition to the MAP kinase pathway (Corrales et al., 2005). Forskolin, an adenylyl cyclase agonist, stimulates cAMP production and hence activates signalling downstream of PKA (cAMP-dependent protein kinase). Cells treated with KCl and forskolin together caused TLK2 to translocate from the nucleus to the cytoplasm as visualised by immunofluorescence (Figure 5C) and quantified by log2(nucleus/cytoplasm) of TLK2 intensity (Figure 5D). These data suggest the nucleocytoplasmic shuttling of TLK2 that occurs during neuronal differentiation might also be regulated in neurons by activity-dependent signalling.

## Discussion

We have undertaken the first characterisation of neuronal TLK2 expression and localisation. We found that TLK2 is expressed in the hippocampus and cerebellum, brain regions that are affected in neurodevelopmental disorders linked with disrupted motor function and cognition. In addition to canonical full length TLK2, we observed splice diversity at the N-terminus of the protein, including variants that lack an NLS. Furthermore, cytoplasmic TLK2 increased during neuronal differentiation, driven by both enhanced expression of TLK2 variants lacking the NLS, and nuclear export of full length TLK2. We also found that the localisation of TLK2 can be regulated acutely by mimicking synaptic activity. Our data suggest future studies into the pathological mechanisms of MRD57 should consider cytoplasmic TLK2 signalling in developing and mature neurons.

### Expression and alternative splicing of neuronal TLK2

Our analysis of TLK2 transcripts in mouse brain, SH-SY5Y cells, and publicly available RNAseq datasets revealed broad conservation of full length TLK2 and shorter transcripts lacking N-terminal exons. Interestingly the canonical, and longest, human Ensembl isoform, TLK2-201, comprising 22 exons is weakly expressed, except in aorta and testis (Figure S1), while the most abundant variant in brain and across the body is TLK-202, which lacks exons 5 and 12. Indeed the other major isoforms that are expressed (TLK-203 and TLK-213) also lack exon 12. TLK-213 is the major annotated short isoform that lacks the N-terminus and NLS and is the human homologue of the short mTLK2-207 isoform that was first identified in mouse testis (Shalom and Don, 1999). However, when we assessed the splice diversity of TLK2 transcripts in SH-SY5Y cells during neuronal differentiation we failed to observe TLK2-213, but instead discovered several novel isoforms that lack the NLS encoded by exon 3.

These variants were only weakly detected in HEK293 cDNA, suggesting they might be upregulated in neurons. When we immunoblotted TLK2 in SH-SY5Y cells using an antibody with a C-terminal epitope, we observed bands that broadly correspond to long and short isoforms. However, due to the minor differences in molecular mass arising from the various splicing events it is not possible to correlate specific protein bands with specific splice variants.

Interestingly, human TLK1, which shares 78% homology with full length TLK2, also possesses a short isoform, TLK1B, lacking the N-terminus and hence the NLS. Constitutive expression of TLK1B is highest in testis, liver and lung, but low in brain (Li et al., 2001). Furthermore, TLK1B expression can be induced by genotoxic stress, such as ionising radiation, through translational regulation of an alternative start site (Li et al., 2001). Surprisingly, in contrast to short TLK2, TLK1B localises to the nucleus where it coordinates DNA repair, even in the absence of the TLK1 NLS (Li et al., 2001; Sunavala-Dossabhoy et al., 2005), suggesting TLK1B is targeted to the nucleus by an alternative mechanism.

Using splice junction-specific *in situ* probes, we were able to examine the relative distributions of long and short TLK2 isoforms in the mouse brain. TLK2 was expressed in the major cell types of the hippocampus and cerebellum, although it was notable that expression in excitatory neurons was easily observed, but inhibitory interneurons were not detectably stained. This is in contradiction to single cell RNAseq analyses of mouse hippocampus and cerebellum, which detected abundant TLK2 transcripts in various interneuron cell types (Habib et al., 2016; Kozareva et al., 2021). We observed differential expression of TLK2 isoforms in excitatory cell types of the hippocampal formation, with intriguing differences between CA1 and CA3 intensity (mTLK2-204) and the blades that comprise the dentate gyrus (mTLK2-201). It is not surprising that there are transcriptional differences between hippocampal cell types, which presumably define their function. However, a deeper understanding of the role of TLK2 in the developing and mature brain is needed to predict the significance of these data.

### Neuronal differentiation- and activity-dependent nucleocytoplasmic shuttling of TLK2

We observed a striking loss of nuclear TLK2 staining in SH-SY5Y cells during neuronal differentiation. Since we did not observe a significant concomitant reduction in protein expression of longer TLK2 isoforms, we concluded that TLK2 was shuttling from the nucleus to the cytoplasm during differentiation. The upregulation of short TLK2 isoform expression we observed is also likely to contribute to the enhancement of cytoplasmic TLK2. This was validated by creating stable SK-N-SH cells cell lines expressing an N-terminally truncated TLK2 variant, which was confined to the cytoplasm. These data agree with the non-nuclear localisation of a short TLK2 variant that was overexpressed in AD293 cells (Mortuza et al., 2018).

Our observation of long TLK2 export from the nucleus during differentiation is consistent with previous data on the cytoplasmic and nuclear distribution of TLK2. Using a C-terminal TLK2 monoclonal antibody, Zhang et al (1999) demonstrated cell cycle-dependent shuttling of TLK2 in cultures of synchronised 3T3 cells. During G1 phase, TLK2 was predominantly outside the nucleus and co-localised with vimentin, an intermediate filament protein. With the onset of S-phase TLK2 was observed to localise to the nuclear periphery and then within the nucleus during late G2 (Zhang et al., 1999). Thus, the nuclear export and cytoplasmic accumulation of TLK2 during neuronal differentiation correlates with these cells exiting the cell cycle. Mechanistically, Zhang et al. attributed the nucleocytoplasmic shuttling of TLK2 to an interaction with 14-3-3 proteins, with binding preventing TLK2 nuclear import. They identified a putative 14-3-3 interaction motif in the C-terminus (amino acids 747-753). Serine phosphorylation of 14-3-3 motifs is required for binding, but the phosphorylation of that motif in TLK2 has not been characterised. Since we observed translocation of TLK2 in response to KCl-induced depolarisation and FSK, it is possible that phosphorylation of TLK2 by a Ca^2+^- and/or cAMP-dependent kinase facilitates 14-3-3 binding and hence retains it in the cytoplasm. There are examples in the literature of transcription factors that require neuronal activity-dependent Ca^2+^ and cAMP signalling for nucleocytoplasmic shuttling, such as the transcriptional activator CRTC1 (Ch’ng et al., 2012). The functions of non-nuclear TLK2 have not been investigated, although its colocalisation with vimentin might suggest regulation of the cytoskeleton (Zhang et al., 1999). This is in-line with the Tousled homologue in *Drosophila*, which has been shown to genetically interact with *rho1* (Gregory et al., 2007), a cytoskeletal regulator, and the *tlk* mutant has altered microtubule and actin filament density in developing follicle cells (Yeh et al., 2015). Other cytoplasmic TLK2 interactors are lacking. Previous TLK2 immunoprecipitation or proximal interactome studies have been conducted in non-neuronal proliferating cell lines, where it might be expected that TLK2 is predominantly nuclear, and these have tended to identify nuclear binding partners (Pavinato et al., 2022; Segura-Bayona et al., 2017).

### The implications of neuronal TLK2 splice variants for MRD57

Our study raises mechanistic questions about the pathology of the neurodevelopmental disorder MRD57, in which patients are haploinsufficient for TLK2, predominantly through *de novo* mutations (Lelieveld et al., 2016; Pavinato et al., 2022; Reijnders et al., 2018; Woods et al., 2022). The important role of TLK2 in cell cycle regulation and DNA repair has led to the hypothesis that the proliferation and differentiation of neural progenitors is affected in MRD57 patient brain development (Segura-Bayona and Stracker, 2019). MRD57 phenotypes are presumably due to processes involving TLK2 that are not compensated by TLK1, such as non-nuclear functions in post-mitotic neurons. Surprisingly, the TLK2 knockout mouse did not display any cell cycle defects, due to compensation by TLK1, except in the placenta where TLK1 expression is low (Segura-Bayona et al., 2017). However, the knockout mouse might not be a good model for MRD57 as there were no reported brain or behavioural anomalies in the TLK2 conditional heterozygous or homozygous mutant mice (Segura-Bayona et al., 2017).

It is notable that MRD57 patient missense mutations have not been identified in exons 2, 4-7 or 12. This suggests that these regions are either not necessary for TLK2 function, or their mutation is tolerated through compensation by alternative splice variants lacking these exons. Relevant to our study, one patient has been identified with a homozygous substitution mutation (K55E) adjacent to the NLS (Töpf et al., 2020), and it would be interesting to study the localisation and nuclear shuttling of this TLK2 mutant in neurons. Furthermore, future studies of neuronal TLK2 should focus on identifying its neuronal cytoplasmic substrates to advance our understanding of TLK2 in developing and mature neurons and the pathology of MRD57.

## Credit author statement

**Lubna Nuhu-Soso**: Writing – review & editing, Writing – original draft, Visualization, Methodology, Investigation, Formal analysis,

Conceptualization. **Heidi Denton**: Investigation, Methodology, Formal analysis. **Darren L. Goffin**: Supervision, Writing – review & editing, Methodology, Conceptualization. **Ines Hahn**: Writing – review & editing, Supervision, Funding acquisition.

**Gareth J.O. Evans**: Writing – review & editing, Writing – original draft, Visualization, Supervision, Project administration, Methodology, Funding acquisition, Formal analysis, Data curation, Conceptualization.

## Acknowledgements

We are grateful to Prof Betsy Pownall and Dr Harry Isaacs (University of York) for valuable advice regarding the *in situ* hybridisation experiments. We thank the Bioscience Technology Facility at the University of York for providing access to confocal microscopes.

