## Supplementary materials for "Neuronal differentiation and activity drive nucleocytoplasmic shuttling of the intellectual disability kinase TLK2"

**Supplemental Materials (Nuhu-Soso et al.)**

**Figure S1. Human tissue TLK2 transcript distribution.** Heatmap of human TLK2 transcript expression in the indicated tissues, expressed as percent abundance. RNAseq data derived from the GTEx Portal, accession number phs000424.v8.p2. Transcripts highlighted in blue are also included in the table in **Fig 1B**.

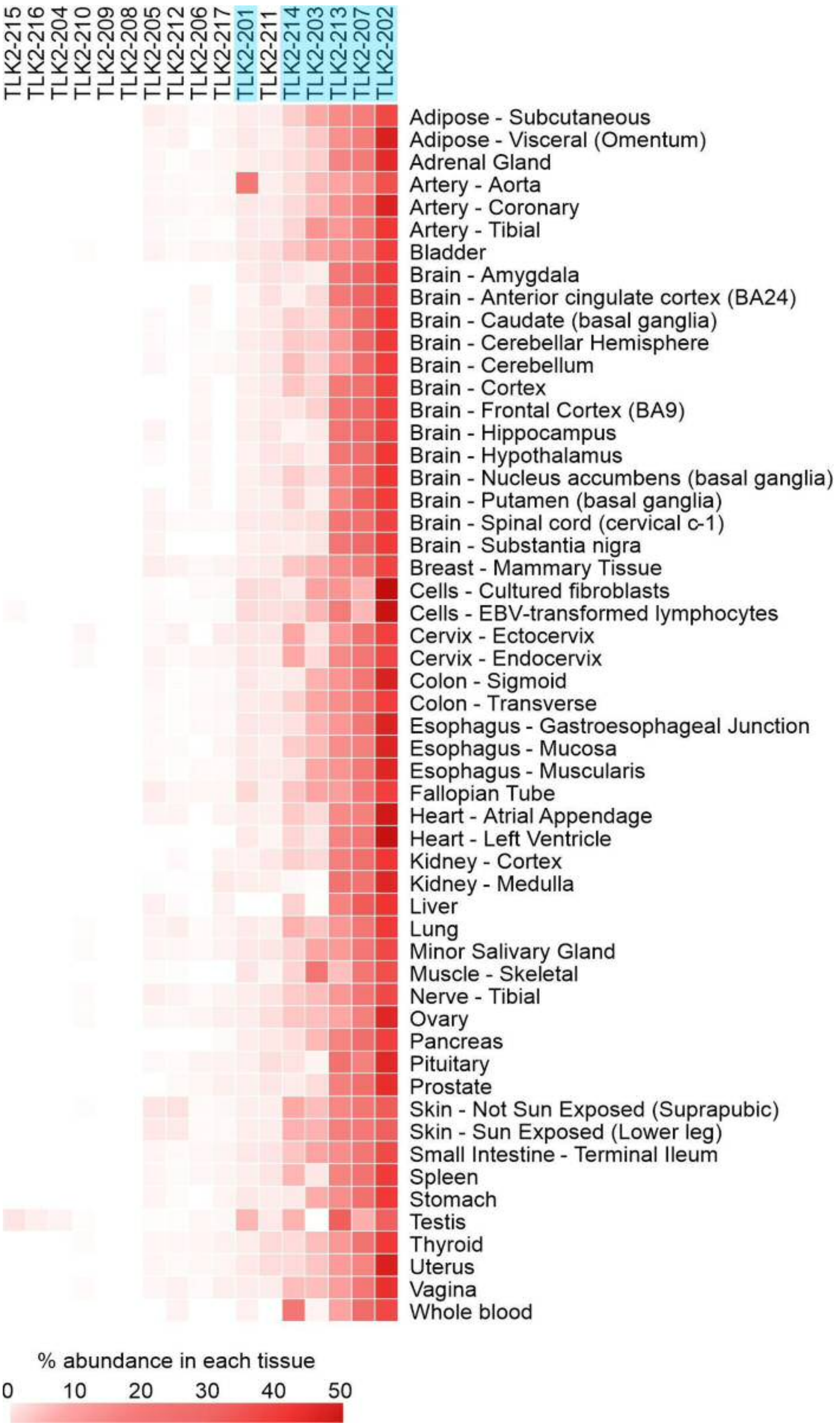

**Figure S2. TLK2 transcripts and their expression in the mouse brain.**
**A.** Exonic structure of mouse TLK2, mapped onto a schematic representation of its protein domains. NLS, nuclear localisation sequence; CC, coiled-coil domain **B.** Table of % TLK2 transcript abundance in the mouse hippocampus and cerebellum derived from HPA RNAseq data plotted in **C**.. Exons labelled in red are predicted to be non-protein coding. Only transcripts present >2% in either brain region are included. **C.** Heatmap of TLK2 transcript expression in the indicated brain regions, expressed as percent abundance. Data derived from HPA RNAseq dataset (Sjöstedt et al., 2020). Transcripts highlighted in blue are included in the table in **B**.

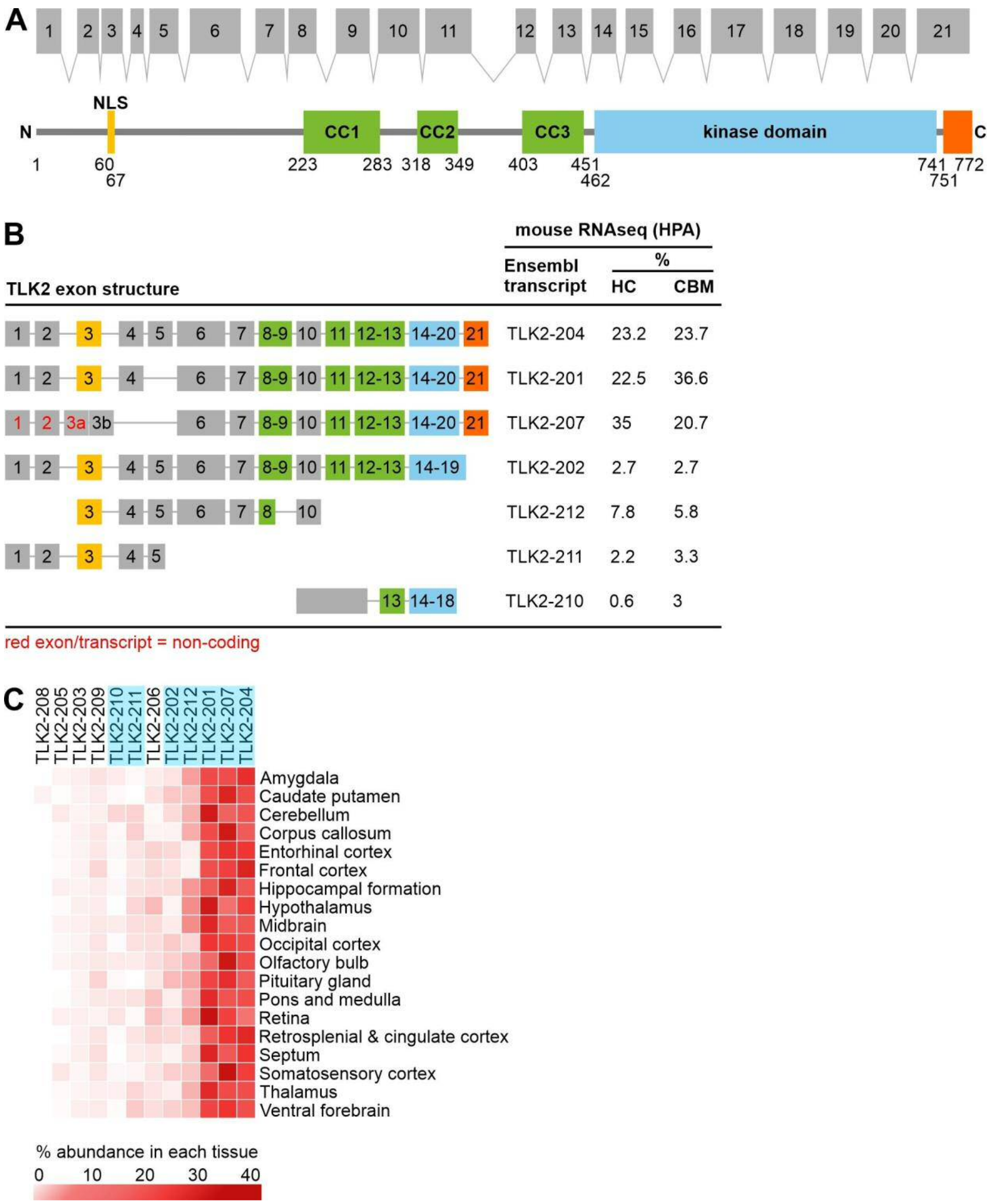

**Figure S3. Sequencing of novel human TLK2 N-terminal splice variants.** cDNA prepared from differentiating SH-SY5Y cells was subjected to PCR with primers located in exons 1 (forward) and 9 (reverse) of human TLK2. The six PCR products obtained (see Fig. 3) were excised from an agarose gel and subjected to Sanger sequencing. The sequences obtained for bands 1,2,4,5 and 6 are presented as an alignment, with each exon indicated by shading in different colours. Band 3 yielded a mix of sequences suggesting multiple transcripts of the same molecular mass. We identified three known transcripts (TLK2-209, -227 and -230) with very similar masses to band 3 within exons 1-9 and include an alignment of these.

```
                       Exon 1 (from 31 bp)
TLK2-201/203 GGAGGCAGGAGTTACTGGAGGCCAGGTTCACTGGGGTTGGCGTAAGTAAGGGGCCACTCA
TLK2-202/224 GGAGGCAGGAGTTACTGGAGGCCAGGTTCACTGGGGTTGGCGTAAGTAAGGGGCCACTCA
Novel a      GGAGGCAGGAGTTACTGGAGGCCAGGTTCACTGGGGTTGGCGTAAGTAAG----------
Novel b      GGAGGCAGGAGTTACTGGAGGCCAGGTTCACTGGGGTTGGCGTAAGTAAGGGGCCACTCA
Novel c      GGAGGCAGGAGTTACTGGAGGCCAGGTTCACTGGGGTTGGCGTAAGTAAG----------

                              Exon 2
TLK2-201/203 ACAGTGAGTCTTCCAACCAGAGTTTGTGCAGTGTGGGGTCCTTGAGTGATAAAGAAGTAG
TLK2-202/224 ACAGTGAGTCTTCCAACCAGAGTTTGTGCAGTGTGGGGTCCTTGAGTGATAAAGAAGTAG
Novel a      ------------------------------------------------------------
Novel b      ACAGTGAGTCTTCCAACCAGAGTTTGTGCAGTGTGGGGTCCTTGAGTGATAAAGAAGTAG
Novel c      ------------------------------------------------------------

                              Exon 3
TLK2-201/203 AGACTCCTGAGAAAAAGCAGAATGACCAGCGAAATCGGAAAAGAAAAGCTGAACCATATG
TLK2-202/224 AGACTCCTGAGAAAAAGCAGAATGACCAGCGAAATCGGAAAAGAAAAGCTGAACCATATG
Novel a      ------------------------------------------------------------
Novel b      AG----------------------------------------------------------
Novel c      ------------------------------------------------------------

                              Exon 4
TLK2-201/203 AAACTAGCCAAGGGAAAGGCACTCCTAGGGGACATAAAATTAGTGATTACTTTGAGTTTG
TLK2-202/224 AAACTAGCCAAGGGAAAGGCACTCCTAGGGGACATAAAATTAGTGATTACTTTGAG----
Novel a      ------------------------------------------------------------
Novel b      ------------------------------------------------------------
Novel c      ------------------------------------------------------------

                              Exon 5
TLK2-201/203 CTGGGGGAAGCGGGCCAGGAACCAGCCCTGGCAGAAGTGTTCCACCAGTTGCACGATCCT
TLK2-202/224 ------------------------------------------------------------
Novel a      ------------------------------------------------------------
Novel b      ------------------------------------------------------------
Novel c      ------------------------------------------------------------


TLK2-201/203 CACCGCAACATTCCTTATCCAATCCCTTACCGCGTCGAGTAGAACAGCCTCTCTATGGTT
TLK2-202/224 --------------------------------CGTCGAGTAGAACAGCCTCTCTATGGTT
Novel a      --------------------------------CGTCGAGTAGAACAGCCTCTCTATGGTT
Novel b      --------------------------------CGTCGAGTAGAACAGCCTCTCTATGGTT
Novel c      --------------------------------CGTCGAGTAGAACAGCCTCTCTATGGTT
```

```
                                 Exon 6
TLK2-201/203 TAGATGGCAGTACTGCAAAGGAGGCCTCAGAAGAGCAGTCTGCCCTGCCAACCCTCATGT
TLK2-202/224 TAGATGGCAGTACTGCAAAGGAGGCCTCAGAAGAGCAGTCTGCCCTGCCAACCCTCATGT
Novel a      TAGATGGCAGTACTGCAAAGGAGGCCTCAGAAGAGCAGTCTGCCCTGCCAACCCTCATGT
Novel b      TAGATGGCAGTACTGCAAAGGAGGCCTCAGAAGAGCAGTCTGCCCTGCCAACCCTCATGT
Novel c      TAGATGGCAGTACTGCAAAGGAGGCCTCAGAAGAGCAGTCTGCCCTGCCAACCCTCATGT

TLK2-201/203 CAGTGATGTTAGCAAAACCTCGACTTGACACAGAGCAGTTAGCGCCAAGGGGAGCTGGCC
TLK2-202/224 CAGTGATGTTAGCAAAACCTCGACTTGACACAGAGCAGTTAGCGCCAAGGGGAGCTGGCC
Novel a      CAGTGATGTTAGCAAAACCTCGACTTGACACAGAGCAGTTAGCGCCAAGGGGAGCTGGCC
Novel b      CAGTGATGTTAGCAAAACCTCGACTTGACACAGAGCAGTTAGCGCCAAGGGGAGCTGGCC
Novel c      CAGTGATGTTAGCAAAACCTCGACTTGACACAGAGCAGTTAGCGCCAAGGGGAGCTGGCC

                                             Exon 7
TLK2-201/203 TCTGCTTCACTTTCGTCTCTGCTCAACAAAACAGCCCTTCGTCCACGGGGTCTGGCAATA
TLK2-202/224 TCTGCTTCACTTTCGTCTCTGCTCAACAAAACAGCCCTTCGTCCACGGGGTCTGGCAATA
Novel a      TCTGCTTCACTTTCGTCTCTGCTCAACAAAACAGCCCTTCGTCCACGGGGTCTGGCAATA
Novel b      TCTGCTTCACTTTCGTCTCT----------------------------------------
Novel c      TCTGCTTCACTTTCGTCTCT----------------------------------------

TLK2-201/203 CAGAACATTCTTGCAGCTCCCAGAAACAGATCTCCATCCAGCACAGGCAGACCCAGTCTG
TLK2-202/224 CAGAACATTCTTGCAGCTCCCAGAAACAGATCTCCATCCAGCACAGGCAGACCCAGTCTG
Novel a      CAGAACATTCTTGCAGCTCCCAGAAACAGATCTCCATCCAGCACAGGCAGACCCAGTCTG
Novel b      ------------------------------------------------------------
Novel c      ------------------------------------------------------------

                                 Exon 8
TLK2-201/203 ACCTCACAATAGAAAAAATATCTGCACTAGAAAACAGTAAGAACTCTGACTTAGAGAAGA
TLK2-202/224 ACCTCACAATAGAAAAAATATCTGCACTAGAAAACAGTAAGAACTCTGACTTAGAGAAGA
Novel a      ACCTCACAATAGAAAAAATATCTGCACTAGAAAACAGTAAGAACTCTGACTTAGAGAAGA
Novel b      ------------------------------------------------------------
Novel c      ------------------------------------------------------------

                                                  Exon 9
TLK2-201/203 AGGAAGGAAGAATAGATGATTTATTAAGAGCCAACTGTGATTTGAGACGACAGATAGATG
TLK2-202/224 AGGAAGGAAGAATAGATGATTTATTAAGAGCCAACTGTGATTTGAGACGACAGATAGATG
Novel a      AGGAAGGAAGAATAGATGATTTATTAAGAGCCAACTGTGATTTGAGACGACAGATAGATG
Novel b      -----------------------------GCCAACTGTGATTTGAGACGACAGATAGATG
Novel c      -----------------------------GCCAACTGTGATTTGAGACGACAGATAGATG

TLK2-201/203 AACAGCAAAAGATGCTAGAGA
TLK2-202/224 AACAGCAAAAGATGCTAGAGA
Novel a      AACAGCAAAAGATGCTAGAGA
Novel b      AACAGCAAAAGATGCTAGAGA
Novel c      AACAGCAAAAGATGCTAGAGA
```

**Options for Band 3 - using Band 1 sequence**

```
                                 Exon 1 (from 31 bp)
```

```
TLK2-209      GGAGGCAGGAGTTACTGGAGGCCAGGTTCACTGGGGTTGGCGTAAGTAAGGGGCCACTCA
TLK2-227      GGAGGCAGGAGTTACTGGAGGCCAGGTTCACTGGGGTTGGCGTAAGTAAGGGGCCACTCA
TLK2-230      GGAGGCAGGAGTTACTGGAGGCCAGGTTCACTGGGGTTGGCGTAAGTAAGGGGCCACTCA

                                     Exon 2
TLK2-209      ACAGTGAGTCTTCCAACCAGAGTTTGTGCAGTGTGGGGTCCTTGAGTGATAAAGAAGTAG
TLK2-227      ACAGTGAGTCTTCCAACCAGAGTTTGTGCAGTGTGGGGTCCTTGAGTGATAAAGAAGTAG
TLK2-230      ACAGTGAGTCTTCCAACCAGAGTTTGTGCAGTGTGGGGTCCTTGAGTGATAAAGAAGTAG

                                     Exon 3
TLK2-209      AGACTCCTGAGAAAAAGCAGAATGACCAGCGAAATCGGAAAAGAAAAGCTGAACCATATG
TLK2-227      AGACTCCTGAGAAAAAGCAGAATGACCAGCGAAATCGGAAAAGAAAAGCTGAACCATATG
TLK2-230      AGACTCCTGAGAAAAAGCAGAATGACCAGCGAAATCGGAAAAGAAAAGCTGAACCATATG

                                     Exon 4
TLK2-209      AAACTAGCCAAGGGAAAGGCACTCCTAGGGGACATAAAATTAGTGATTACTTTGAG----
TLK2-227      AAACTAGCCAAGGGAAAGGCACTCCTAGGGGACATAAAATTAGTGATTACTTTGAG----
TLK2-230      AAACTAGCCAAGGGAAAGGCACTCCTAGGGGACATAAAATTAGTGATTACTTTGAGTTTG

                                     Exon 5
TLK2-209      ------------------------------------------------------------
TLK2-227      ------------------------------------------------------------
TLK2-230      CTGGGGGAAGCGGGCCAGGAACCAGCCCTGGCAGAAGTGTTCCACCAGTTGCACGATCCT


TLK2-209      --------------------------------CGTCGAGTAGAACAGCCTCTCTATGGTT
TLK2-227      --------------------------------CGTCGAGTAGAACAGCCTCTCTATGGTT
TLK2-230      CACCGCAACATTCCTTATCCAATCCCTTACCGCGTCGAGTAGAACAGCCTCTCTATGGTT

                                     Exon 6
TLK2-209      TAGATGGCAGTACTGCAAAGGAGGCCTCAGAAGAGCAGTCTGCCCTGCCAACCCTCATGT
TLK2-227      TAGATGGCAGTACTGCAAAGGAGGCCTCAGAAGAGCAGTCTGCCCTGCCAACCCTCATGT
TLK2-230      TAGATGGCAGTACTGCAAAGGAGGCCTCAGAAGAGCAGTCTGCCCTGCCAACCCTCATGT


TLK2-209      CAGTGATGTTAGCAAAACCTCGACTTGACACAGAGCAGTTAGCGCCAAGGGGAGCTGGCC
TLK2-227      CAGTGATGTTAGCAAAACCTCGACTTGACACAGAGCAGTTAGCGCCAAGGGGAGCTGGCC
TLK2-230      CAGTGATGTTAGCAAAACCTCGACTTGACACAGAGCAGTTAGCGCCAAGGGGAGCTGGCC

                                                 Exon 7
TLK2-209      TCTGCTTCACTTTCGTCTCT----------------------------------------
TLK2-227      TCTGCTTCACTTTCGTCTCTGCTCAACAAAACAGCCCTTCGTCCACGGGGTCTGGCAATA
TLK2-230      TCTGCTTCACTTTCGTCTCT----------------------------------------

TLK2-209      --------------------------------------------------------TCTG
TLK2-227      CAGAACATTCTTGCAGCTCCCAGAAACAGATCTCCATCCAGCACAGGCAGACCCAG----
TLK2-230      ------------------------------------------------------------

                                     Exon 8
TLK2-209      ACCTCACAATAGAAAAAATATCTGCACTAGAAAACAGTAAGAACTCTGACTTAGAGAAGA
TLK2-227      ------------------------------------------------------------
TLK2-230      ------------------------------------------------------------
```

```
                                                     Exon 9
TLK2-209     AGGAAGGAAGAATAGATGATTTATTAAGAGCCAACTGTGATTTGAGACGACAGATAGATG
TLK2-227     ----------------------------GCCAACTGTGATTTGAGACGACAGATAGATG
TLK2-230     ----------------------------GCCAACTGTGATTTGAGACGACAGATAGATG

TLK2-209     AACAGCAAAAGATGCTAGAGA
TLK2-227     AACAGCAAAAGATGCTAGAGA
TLK2-230     AACAGCAAAAGATGCTAGAGA
```

**Raw images of gels presented in figures.**

### Figure 2

Gel-doc DNA image with crops for 2A and 2B

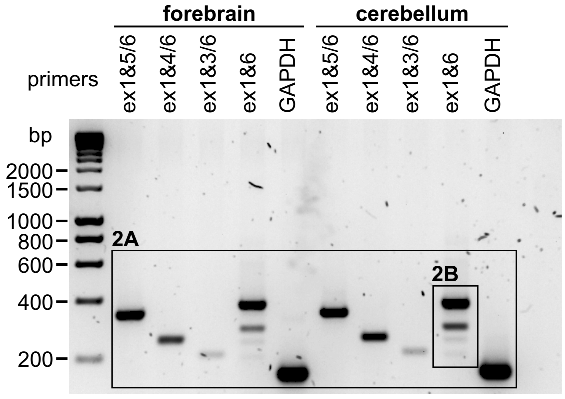


### Figure 3A

Gel-doc DNA image with crops

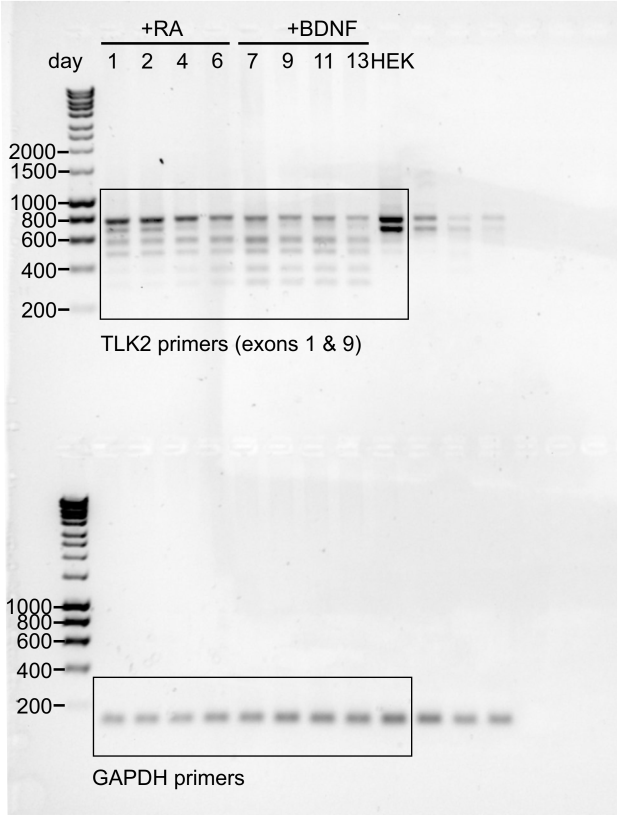


Gel-doc DNA image with crop

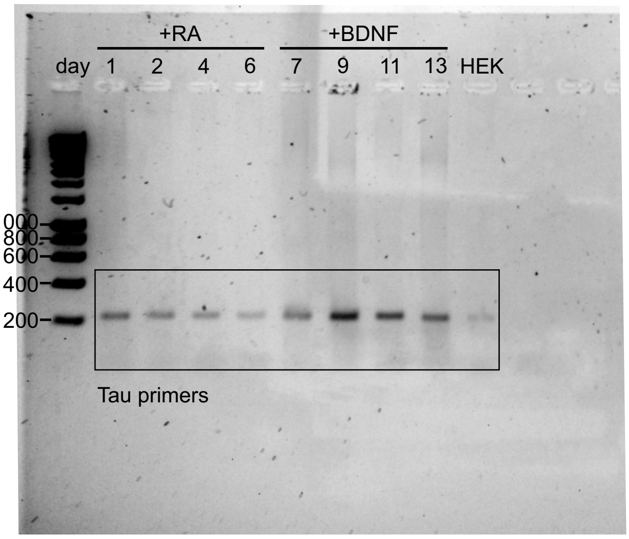

**Figure 4A**

anti-TLK2 - membrane image

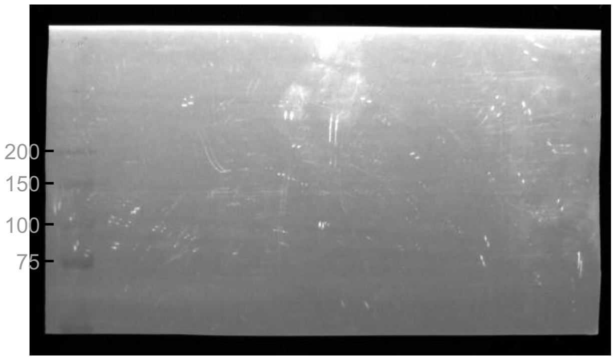


anti-TLK2 - iBright image with crop

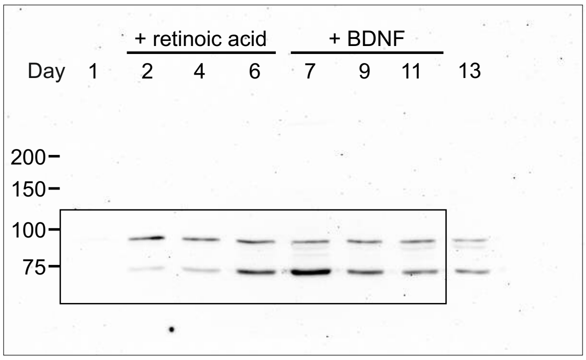


anti-actin - membrane image

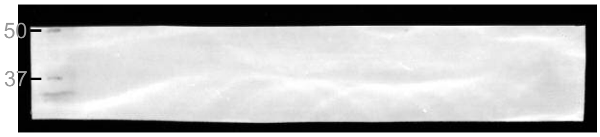


anti-actin - iBright image with crop

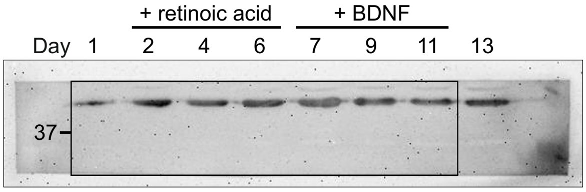


anti-β3-tubulin - membrane image

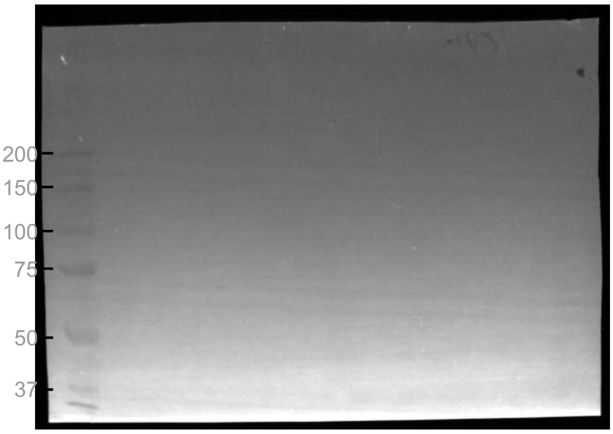


anti-β3-tubulin - iBright image with crop

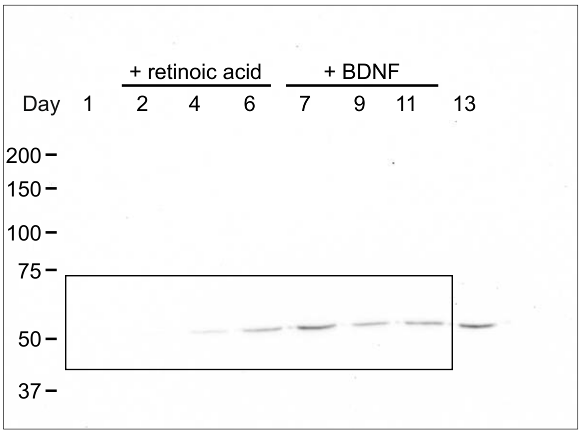

### Figure 5B - top blots (TLK2-202)

#### anti-FLAG - membrane image

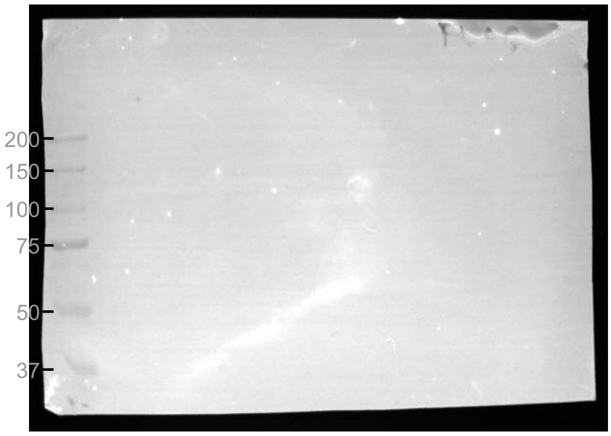


#### anti-FLAG - iBright image with crop

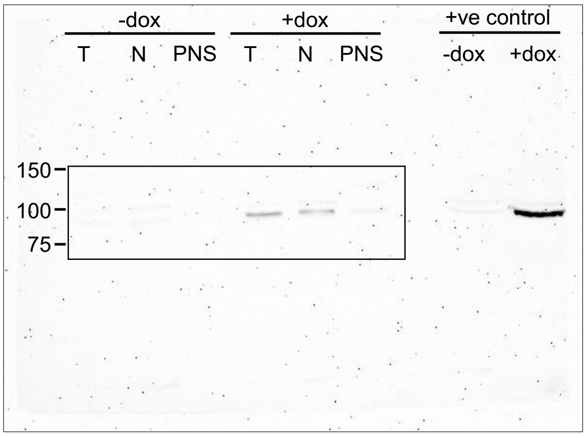


#### anti-GAPDH - membrane image

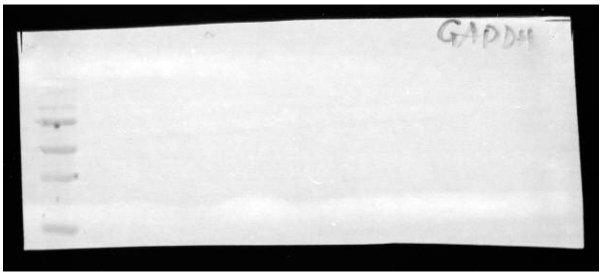

#### anti-GAPDH - iBright image with crop

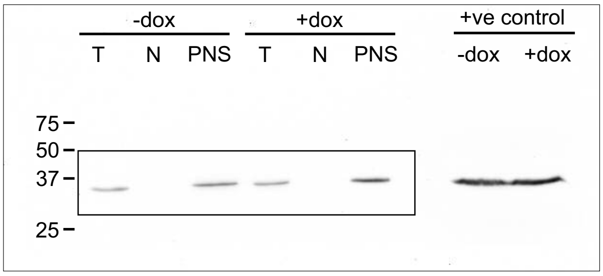


#### anti-histone H3 - membrane image

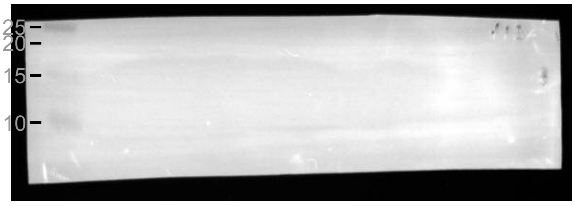


#### anti-histone H3 - iBright image with crop

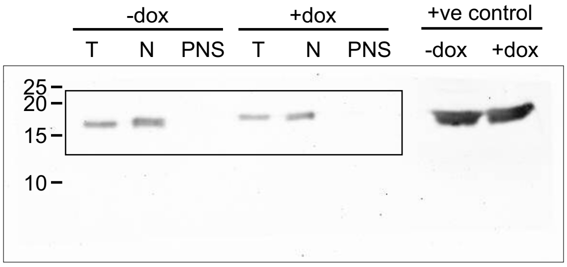

**Figure 5B - bottom blots (TLK2-213)**

anti-FLAG - membrane image

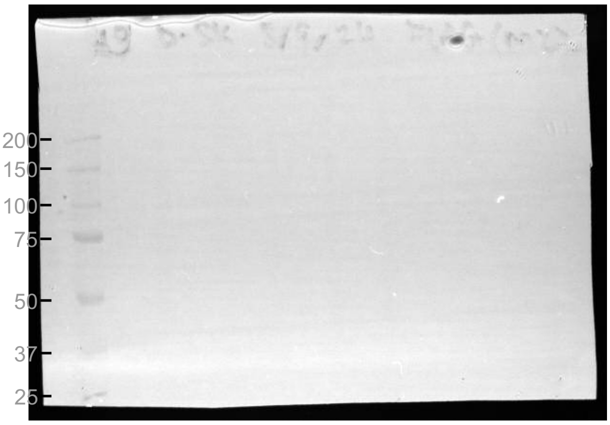


anti-FLAG - iBright image with crop

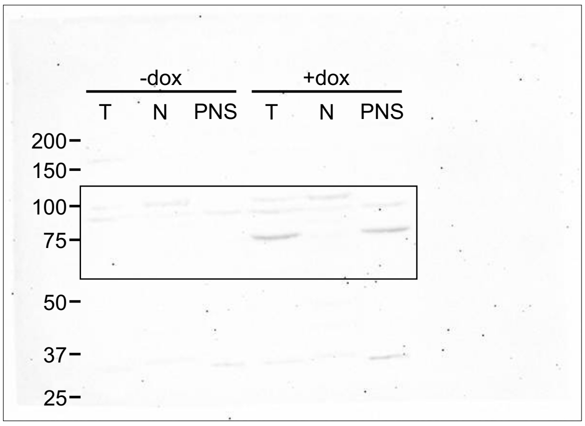


anti-GAPDH - membrane image

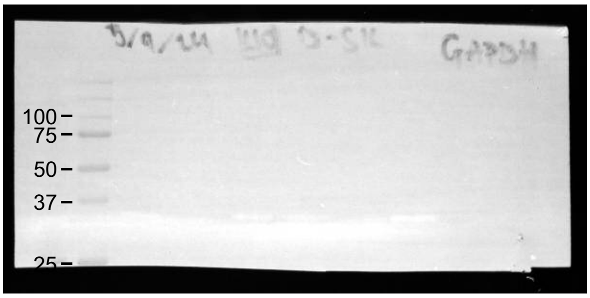


anti-GAPDH - iBright image with crop

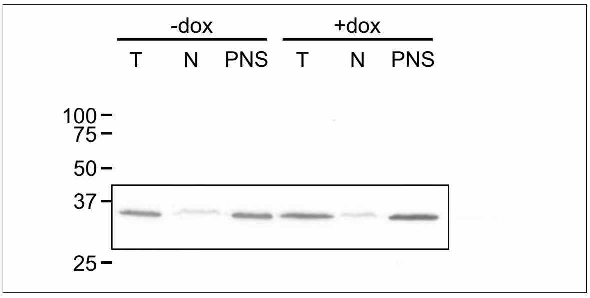


anti-histone H3 - membrane image

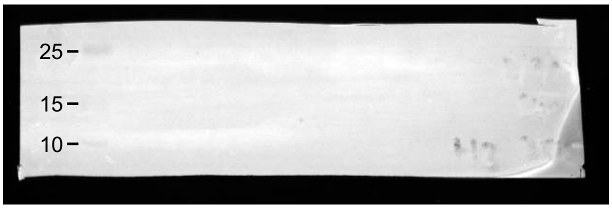


anti-histone H3 - iBright image with crop

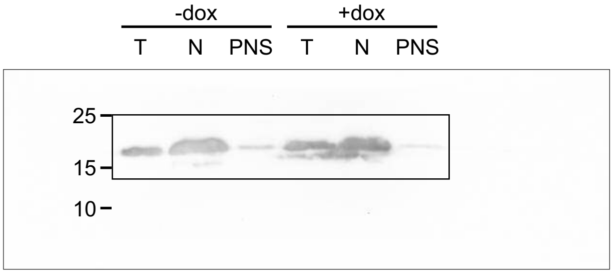
